# All-Optical Multimodal Mapping of Single Cell-Type–Specific Metabolic Activities via REDCAT

**DOI:** 10.1101/2024.11.07.622511

**Authors:** Yajuan Li, Zhaojun Zhang, Archibald Enninful, Negin Farzad, Presha Rajbhandari, Jungmin Nam, Xiaoyu Qin, Jorge Villazon, Anthony A. Fung, Hongje Jang, Zhiliang Bai, Nancy R. Zhang, Brent R. Stockwell, Rong Fan, Mina L. Xu, Zongming Ma, Lingyan Shi

## Abstract

Metabolism underlies cell growth, survival, and function, yet its activities vary widely across cell types and tissue environments. Spatially resolving these processes *in situ* at single-cell resolution is essential to advance our understanding of cellular function and tissue physiology in health and disease. However, existing approaches are limited by either destructive workflows, insufficient spatial resolution and biochemical specificity, or lack of direct linkage to cell identity. Here, we present Raman Enhanced Delineation of Cell Atlases in Tissues (REDCAT), a multimodal all-optical platform that integrates stimulated Raman scattering, autofluorescence redox imaging, second harmonic generation, and high-plex immunofluorescence to co-map metabolic activities and cell types within the same tissue section. REDCAT achieves subcellular resolution profiling of protein, lipid, redox, and nuclear acid metabolism, together with extracellular matrix composition, in both FFPE and fresh-frozen human tissues. Applied to normal lymph nodes, REDCAT delineated distinct redox and lipid remodeling programs across germinal center B-cell zones and immune subsets, highlighting cell-type–specific metabolic specialization. In lymphoma, it revealed profound metabolic reprogramming, including extensive lipid accumulation, nuclear metabolic heterogeneity, and a transitional metabolic state associated with transformation from chronic lymphocytic leukemia to diffuse large B-cell lymphoma, thereby illuminating tumor evolution *in situ*. In human liver, REDCAT resolved cell-type–specific lipid droplet diversity and zonation-dependent nuclear metabolic gradients, uncovering new principles of spatial metabolic organization. By directly linking cell identity with spatial metabolic states at single-cell or subcellular resolution, REDCAT establishes a broadly applicable framework for studying immune function, tumor progression, and tissue physiology, and offers a new path to deciphering the metabolic basis of health and disease.

## Introduction

Metabolism is a fundamental process that profoundly influences cellular function in normal physiological states and across diverse disease conditions, including immune responses and cancer progression ^1–4^. Metabolic activity encompasses a collection of chemical reactions and processes that occur within a cell or organism to sustain life, generate energy, and mediate the synthesis and breakdown of biomolecules ^1–3,5^. It reflects how actively a cell is processing nutrients, synthesizes biomolecules, and maintains redox and energy balance ^2,6,7^. Metabolic activity within a cell is dynamically controlled by a combination of intrinsic programs and extrinsic cues, allowing the cell to adapt to its functional state and microenvironmental context ^8–10^. Characterizing the metabolic activity of individual cells *in situ* is fundamental to improving our understanding of complex biological systems. Despite recent advances in spatial omics ^11–18^ that have revealed single-cell transcriptional or protein profiles that can potentially infer metabolic processes, heterogeneity, and interaction between spatial neighborhoods within a tissue, these approaches neither directly capture the dynamic state of cellular metabolism nor establish functional links between metabolic activity and specific cell types or states.

The emerging field of spatial metabolomics aims to profile metabolic fingerprints of tissues at different pathophysiological states ^14,19^. Progress in this field relies heavily on technological innovations that continually expand our capability to detect and quantify the thousands of metabolites present in biological samples ^6,20–22^. A wide range of mass spectrometry imaging (MSI) methods ^22–26^ including Matrix-Assisted Laser Desorption/Ionization (MALDI) ^27^, Desorption Electrospray Ionization (DESI) ^23^, and Secondary Ion Mass Spectrometry (SIMS), have enabled researchers to gain spatial information about molecular distributions within biological samples. When these approaches are integrated with multiplexed protein markers, they offer the capability to characterize metabolic cellular heterogeneity with high sensitivity, spatial resolution, and chemical coverage. For example, a recent method named SpaceM, which combines fluorescence imaging with MALDI MSI ^28,29^ has been employed to map the metabolic profiles of single cells. Spatially resolved metabolomic profiling methods based on TOF-SIMS ^30^, such as 3D-SMF ^31^ and scSpaMet ^21^, have demonstrated spatial mapping of proteins and metabolites in tissues. However, the fundamental limitation of most MSI techniques includes the trade-off between molecular coverage and spatial resolution as well as the inherently destructive nature, which restricts their application in studies requiring longitudinal measurements or across different modalities on the same tissue section. This destructive aspect complicates truly integrated multi-omics analyses, as it poses challenges for precise co-registration and direct correlation of metabolic data with other molecular or cellular data — such as multiplex protein-based cell typing or transcriptomic profiling — within the same cells or even subcellular regions.

Raman imaging offers advantages because it is not affected by water interference, making it well-suited for biological samples and enabling chemical or metabolic imaging at subcellular resolution in aqueous environments ^32–40^. For example, Raman spectroscopy can characterize the biomolecular composition of clinical histological samples and resolve the spatial distribution of metabolites, such as fumarate and cytochrome c, within cells and tissues ^41,42^. Furthermore, Raman microscopy allows for totally nondestructive detection or even tracking of structurally diverse lipid species to dissect their spatiotemporal dynamics *in situ* and can infer single-cell gene expression profiles in live cells ^22,43–45^. However, a major limitation of spontaneous Raman imaging is its slow acquisition speed, attributed to the intrinsically weak Raman scattering cross-section. This challenge is largely overcome by nonlinear Raman techniques, such as stimulated Raman scattering (SRS), which significantly enhance imaging sensitivity ^46^. It has been successfully applied to visualize the subcellular distribution and synthesis of lipids, proteins, nucleic acids and other molecules both *in vitro* and *in vivo* ^46–50^. SRS provides a background-free, linear signal that is directly proportional to molecular concentration, enabling reliable quantitative imaging ^51^. Moreover, hyperspectral SRS generates spectra closely resembling those from spontaneous Raman ^52,53^, making it particularly advantageous for hyperspectrally resolved molecular discrimination, metabolic fingerprinting, and multi-modal integration.

To fill this technological gap, we developed Raman Enhanced Delineation of Cell Atlases in Tissues (REDCAT), an integrated platform for profiling cell-type**–**specific metabolic activities by incorporating SRS probed metabolomics, two-photon excited fluorescence (TPEF), second harmonic generation (SHG) imaging and highly multiplexed immunofluorescence imaging-based cell typing via co-detection by indexing (CODEX). This capability was further enhanced by a computational pipeline using MaxFuse ^54^ to integrate metabolic imaging with CODEX and scRNA-deq data. This multimodal imaging platform enables submicron-resolution measurement of intracellular proteins, lipids, nuclear metabolism, and redox status, all co-registered with a large panel of protein markers used to generate a spatial cell-type map. It also allows for characterizing the heterogeneity of extracellular matrix components, particularly collagen, within the same tissue section. REDCAT is compatible with both fresh-frozen (FF) and formalin-fixed paraffin-embedded (FFPE) tissues from clinical tissue banks. Leveraging this platform, we conducted a systematic analysis of metabolic profiles in human normal and malignant lymphoid tissues, and observed that metabolic redox state, single-nucleus metabolomics, metabolic reprogramming, and lipid droplet (LD) heterogeneity are specific to cell types and spatially regulated in the tissue niches. We further dissected single cell-type**–**specific metabolic state in human liver samples, unravelling metabolic zonation and distance dependent nuclear metabolic heterogeneity from the central vein (CV) to the portal vein (PV). Together, these results have demonstrated REDCAT’s capability to integrate Raman based spatial metabolic imaging with multiplexed protein imaging at single-cell resolution, offering deeper insights into the cell-type**–**specific metabolic activities and metabolic reprogramming in complex tissues.

## Results

### Design and workflow of REDCAT

The workflow of REDCAT is described in Fig. 1. A deparaffinized FFPE or FF tissue section (∼10 µm in thickness) mounted on glass slide was used. To directly link endogenous metabolic states with cell-type**–**specific protein expression on the same tissue section, two-photon excitation fluorescence-integrated stimulated Raman scattering (TPEF-SRS) imaging was first performed to capture label-free metabolic profiles at single-cell resolution. Specifically, the reduced forms of nicotinamide adenine dinucleotide or its phosphorylated variant (NAD(P)H), as well as the oxidized flavin cofactors flavin adenine dinucleotide (FAD) and flavin mononucleotide (FMN) were visualized using TPEF autofluorescence microscopy. The relative abundance of these two classes of metabolic cofactors, defined as the redox ratio, provides a quantitative indicator of cellular redox balance between reduction and oxidation processes. Since NADH and FAD are closely coupled to energy-generating pathways and robustly reflect the bulk redox state, the NADH/FAD ratio is used in this study as a proxy for the cell redox state defined as optical redox ratio ^55^. A higher NADH/FAD ratio reflects a shift toward a more reduced metabolic state, consistent with increased glycolysis and/or impaired oxidative phosphorylation ^56^. Subsequently, label-free SRS imaging was used to acquire signals for total protein, total lipid, unsaturated lipid, and saturated lipid from the same region of interest (ROI), achieved by sequentially tuning the pump laser wavelength to Raman shifts of 2930 cm^−1^, 2850 cm^−1^, 3015 cm^−1^ and 2880 cm^−1^, respectively. The Raman intensity ratio of 2850 cm^−1^ to 2930 cm^−1^ (2850/2930) was calculated to indicate relative lipid abundance, whereas the ratio of 3015 cm^−1^ to 2880 cm^−1^ (3015/2880) reflects the degree of lipid unsaturation, which is associated with membrane fluidity and susceptibility to oxidative stress. In parallel, SHG imaging was applied to detect collagen fibers by exploiting their intrinsic non-centrosymmetric structure, providing complementary information on tissue architecture without the need for exogenous labeling. To further resolve the detailed chemical properties of *in situ* metabolites, hyperspectral SRS images were acquired across 2700-3150 cm^−1^ SRS spectral range, capturing 51 spectral channels at ∼10 cm^−1^ resolution. Label-free redox, SHG, and SRS imaging provide quantitative, non-perturbative, multiplexed, and clinically translatable readouts of metabolism and biomolecular composition, offering advantages over conventional fluorescence imaging, which— although highly sensitive and versatile—relies on exogenous labeling, has limited multiplexing capacity, and is prone to photobleaching. It is worth noting that all images acquired using the TPEF-SRS system were obtained at a uniform spatial resolution of approximately 0.25 µm per pixel, enabling metabolic profiling in subcellular structures, including nuclei and LDs. Lastly, the same tissue section was subsequently mapped using multiplexed immunofluorescence protein imaging via CODEX to identify all major cell types. A panel of ∼40 protein markers was applied, and the imaging was performed at a spatial resolution of roughly 0.5 µm per pixel. Following CODEX imaging, the same tissue section was stained with hematoxylin and eosin (H&E) to enhance visualization of tissue morphology and enable histopathological evaluation by expert pathologists. After all imaging procedures were completed, data from the various modalities were integrated at single-cell resolution on the same tissue section using a custom-designed computational pipeline. Specifically, single cell features from each modality were extracted following cell segmentation, with cell boundaries defined using pretrained Mesmer models. To ensure accurate spatial alignment, affine transformations were applied to register CODEX and TPEF-SRS images at the cellular level, enabling precise association between molecular and metabolic features. Subsequently, MaxFuse pipeline was applied for cross-modal cell matching and joint embedding of all cells in different modalities ^54^. This cross-modality integration empowers the REDCAT platform to generate *in situ* profiles of metabolic redox activity together with protein, lipid, and nuclear metabolic information, quantified cell-by-cell in the tissue section at single-cell resolution. Consequently, cell types determined by canonic protein markers are directly associated with their metabolic states defined by quantitative biochemical indicators including NADH/FAD, lipid/protein, unsaturated/saturated lipid ratios, and comprehensive SRS hyperspectral signatures.

**Fig. 1.**
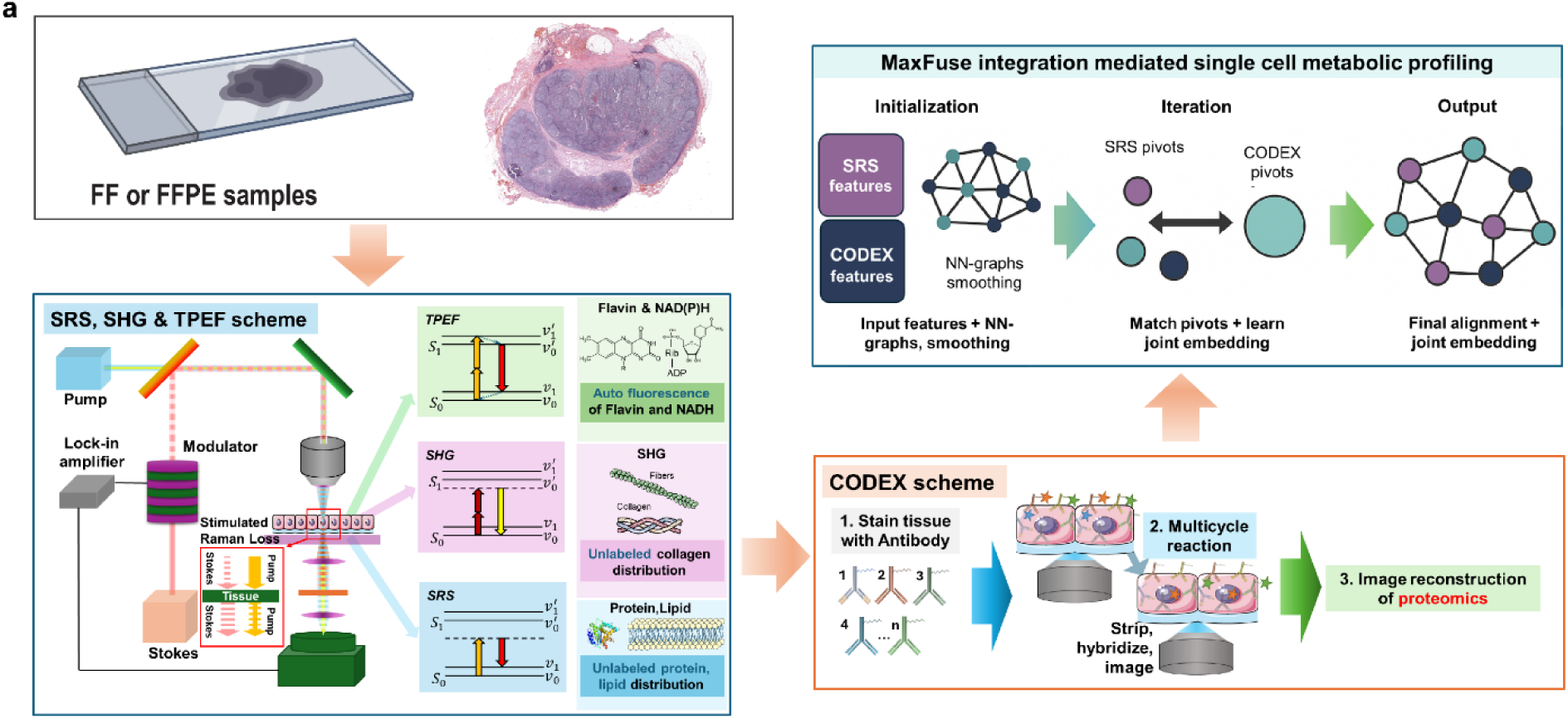
Design and workflow of REDCAT for integrated metabolic and proteomic profiling on the same tissue section. **(a)** Schematic workflow. FF (fresh frozen) or FFPE (formalin-fixed paraffin-embedded) tissue samples are sectioned and placed on a standard glass slide to firstly perform multimodal optical metabolic imaging that combines stimulated Raman scattering (SRS), second harmonic generation (SHG), and two-photon excited fluorescence (TPEF), and the same tissue section is subsequently imaged by highly multiplexed immunofluorescence such as CODEX for spatial protein profiling. Optical metabolic imaging includes autofluorescence for NADH and flavin (redox imaging), SHG for unlabeled collagen distribution, and SRS for label-free protein and lipid mapping. CODEX multiplexed immunofluorescence workflow involves iterative antibody staining, stripping, and imaging, followed by image reconstruction. It routinely images ∼50 protein markers for precise cell tying of highly heterogeneous cell populations in tissue. The resulting metabolic maps of NADH/FAD ratio, lipid/protein ratio, and collagen distribution as well as SRS hyperspectral imaging coupled with CODEX images from the same tissue section were fed into the MaxFuse pipeline. It created a nearest-neighbor (NN) graph for each modality, followed by the smoothing and matching process. Finally, co-registration of SRS and CODEX data enables the mapping of single-cell-resolved and cell-type–specific metabolic profiles.

### REDCAT maps cell-type–specific metabolic activities in healthy human lymph nodes

To demonstrate REDCAT for decoding normal immune physiology, we applied it to a deparaffinized FFPE lymph node tissue section from a healthy donor across a 1.6 mm × 1.6 mm ROI (Extended Fig. 1a). Lymph nodes, as the central component of the secondary lymphoid system, are primarily composed of B and T lymphocytes, organized in coordination with dendritic cells, macrophages, and stromal cells around spatially defined lymphoid follicles ^57–59^. Immune cells within these tissues undergo dynamic processes such as proliferation, differentiation, activation, and responses to extracellular signals or stressors ^57,60^. These processes are tightly regulated by coordinated cell remodeling, niches and interactions accompanied by distinct metabolic reprogramming ^4,58,60,61^.

To spatially investigate single cell-type**–**specific metabolic and molecular features in this immune organ, NADH, FAD, protein, lipid signals were acquired using TPEF-SRS imaging (Extended Fig. 1b), followed by CODEX imaging that was performed on the same tissue section using an antibody panel comprising 48 protein markers (Extended Fig. 2a). This CODEX panel included surface markers for immune cells, cytokine receptors, proliferation markers, and extracellular matrix proteins. Following cell segmentation and unsupervised clustering of single-cell protein expression profiles, we identified nine distinct cell clusters, which were spatially organized in accordance with known histological structures (Fig. 2a, b, Extended Fig. 1a). Specifically, clusters 3, 4, 5, 6 and 8 corresponded to the follicular zone, while clusters 0, 1, 2 and 7 were associated with the interfollicular zone. After image integration, SRS and CODEX images were precisely co-registered to achieve single-cell–level spatial correspondence (Fig. 2c, Extended Fig. 1b). The TPEF-SRS imaging based NADH/FAD, lipid/protein, and unsaturated/saturated lipid ratiometric images were generated (Fig. 2d-f) and the ratio value for each CODEX cluster was extracted. The quantification analysis revealed a positive correlation between the unsaturated/saturated lipid ratio and the NADH/FAD ratio, but a negative correlation between the lipid/protein ratio and either the NADH/FAD or unsaturated/saturated lipid ratios across most clusters (Fig. 2g). These relationships suggest that elevated lipid metabolic activity, particularly through lipid desaturation, is associated with a more reduced cellular redox state. In contrast, total lipid accumulation, as indicated by a high lipid/protein ratio, is associated with a more oxidative redox state. Notably, cells in cluster 4 (yellow in Fig. 2a, b), located in the germinal center (GC) light zone, exhibited a higher NADH/FAD redox ratio and a higher unsaturated to saturated lipid ratio compared to those in cluster 3 (cyan in Fig. 2a, b), which was situated in the dark zone (as marked in Extended Fig. 1a H&E image). Both clusters showed strong expressions of CD20 and CD21, supporting their identification as GC B cells (Fig. 2a). This observation is in line with previous reports showing that GC dark zone B cells (cluster 3) are metabolically oriented toward rapid proliferation and genetic diversification, requiring high energy and biosynthetic flux. These demands lead to increased oxidative stress and a more oxidized redox state. In contrast, light zone (cluster 4) B cells are more metabolically quiescent but engage in antigen processing and presentation. They may upregulate unsaturated lipid remodeling and redox buffering, leading to a more reduced redox state and fluid membranes, to support interactions with T follicular helper (Tfh) cells. In addition to B cells, a distinct population of cells in cluster 8 (green in Fig. 2a, b, h), located between the GC light and dark zones, exhibited a relatively high NADH/FAD ratio (Fig. 2h). These cells expressed CD14, CD38, IFN-γ and podoplanin, suggesting they may represent macrophages/monocytes or dendritic cells actively producing cytokines/chemokines within the GC. Their metabolic profile characterized by a highly reduced redox state indicates active glycolysis (Fig. 2h). Similarly, on the apical side of the same GC follicle, we identified a group of T follicular helper (Tfh) cells expressing CD4, PD-1, CXCR5, and HLA-DR (Fig. 2i). It showed a markedly high level of the NADH/FAD ratio. This cell-type**–**specific glycolysis-associated metabolic phenotype likely reflects the energetic and biosynthetic demands of supporting B cell activation and affinity maturation during the GC reaction.

**Fig.2. Integrated.**
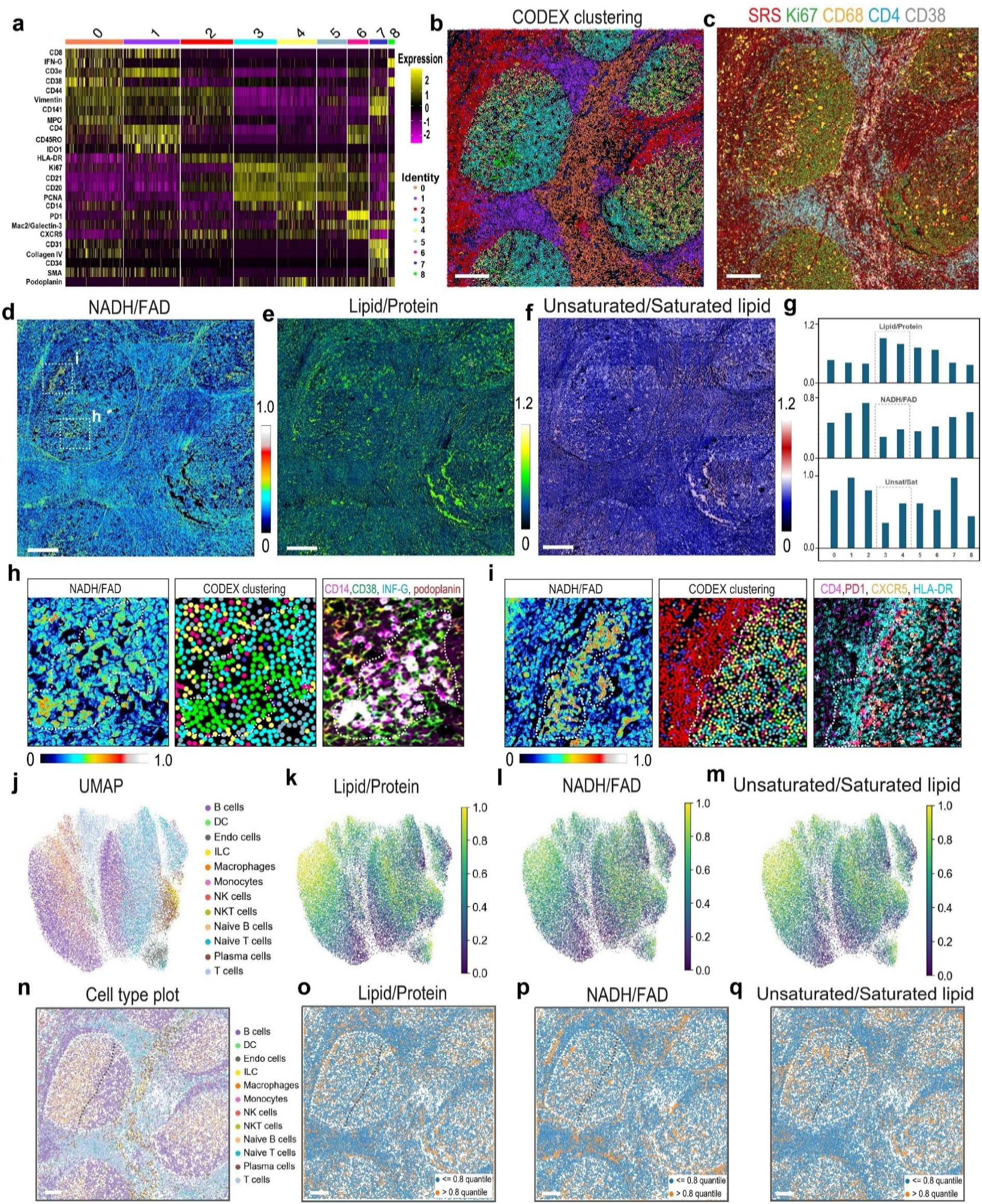
metabolic and cell type profiling of human lymph node tissue via REDCAT. **(a)** Heatmap showing expression levels of protein markers across distinct cell clusters identified by CODEX imaging. Marker expression calibration bar (magenta–yellow) and cluster identities (color dots) are indicated. **(b)** Spatial map of all nine cell clusters from unsupervised clustering of CODEX data after segmentation. Scale bar, 200 μm. **(c)** Representative CODEX image showing expression of Ki67 (green), CD68 (yellow), CD4 (cyan), and CD38 (white), which is further overlaid with the SRS signal (red) of CH3 vibrational modes (total protein signal). Scale bar, 200 μm. **(d–f)** SRS-based metabolic maps of the same tissue section showing (d) NADH/FAD optical redox ratio, (e) lipid/protein ratio, and (f) unsaturated/saturated lipid ratio. Scale bars, 200 μm. **(g)** Quantification of lipid/protein, NADH/FAD, and unsaturated/saturated lipid ratios across the identified cell clusters. Among them, cluster 3 (Cyan in (b)) and cluster 4 (Yellow in (b)) were highlighted. **(h)** Enlarged view of a region from (d) highlighting high NADH/FAD signal (left), corresponding CODEX cluster identities (middle), and immunofluorescence markers for CD14 (magenta), CD38 (green), IFN-γ (cyan), and podoplanin (red) (right). **(i)** Enlarged view of a region from (d) showing high NADH/FAD in a T follicular helper (Tfh) cell-rich area (left), corresponding CODEX clustering (middle), and immunofluorescence for CD4 (magenta), PD-1 (red), CXCR5 (fellow), and HLA-DR (cyan) (right). **(j)** UMAP visualization of single-cell clustering annotated by cell type. **(k–m)** UMAP feature plots showing single-cell lipid/protein ratio (k), NADH/FAD (l), and unsaturated/saturated lipid ratio (m) generated by co-registration of multimodal SRS imaging with CODEX cell typing data. **(n)** Spatial matrix plot displaying cell-type distribution across tissue architecture. Scale bar, 200 μm. **(o–q)** Spatial maps of lipid/protein (o), NADH/FAD (p), and unsaturated/saturated lipid (q) ratios overlaid with cell-type annotations, highlighting metabolic heterogeneity in a follicular zone. Scale bars, 200 μm.

To further evaluate the metabolic activity of each cell type, we annotated individual cells based on protein expression profiles obtained from the CODEX assay (Extended Fig. 1c). While the high-plex protein panel enabled unbiased clustering of tissue pixels into spatially distinct regions and allowed broad classification of major immune and stromal cell types (Extended Fig. 1c, d), it lacked sufficient resolution to discriminate finer immune cell subclasses. In particular, the existing protein markers could not reliably distinguish between naïve, memory, plasma, and GC B cell subsets as well as distinct T cell subsets with specialized functions. To overcome this limitation, we further integrated and co-embedded the CODEX data with a reference single-cell RNA-sequencing (scRNA-seq) atlas from lymph node cells using a computational pipeline MaxFuse (Methods). This multimodal approach provides enhanced subclass-level cell identity and functional annotation, enabling a more detailed interpretation of cell-type**–**specific metabolic features. Using Uniform Manifold Approximation and Projection (UMAP) visualization, we clustered single cells and annotated them into major immune and stromal populations, including subtypes of B cells, T cells, dendritic cells (DCs), monocytes, macrophages, natural killer (NK) cells, endothelial cells, and innate lymphoid cells (ILCs) (Fig. 2j).

We next extracted single-cell lipid/protein ratios, NADH/FAD redox ratios, and unsaturated/saturated lipid ratios quantified from TPEF-SRS imaging and plotted their distributions as histograms (Extended Fig. 1e-g). Cells with ratio values in the top quantile (≥0.8 quartile) were predominantly B and T cells, consistent with their abundance in the lymph node (Extended Fig. 1h-j). To further investigate the relationship between metabolic state and cell identity, we overlaid these metabolic features onto the cell type UMAP. The results revealed a broad distribution of metabolic states within each annotated cell type cluster, including B cells, T cells, monocytes, and macrophages (Fig. 2k-m). This indicates that cell identity alone does not fully define metabolic phenotypes and suggests that context-dependent states such as activation, differentiation, or spatial localization within the tissue may contribute to metabolic reprogramming.

To spatially resolve single-cell metabolic phenotypes across annotated cell types, we reprojected cell types identified by MaxFuse integration of CODEX and snRNA-seq data into their original tissue coordinates using matrix plotting (Fig. 2n). Consistently, we observed that metabolic features (NADH/FAD, lipid/protein, and unsaturated/saturated lipid ratio) did not uniformly correspond to specific cell types (Fig. 2o-q). Nevertheless, functionally relevant spatial patterns emerged. Within GC, most B cells in the light zone exhibit elevated NADH/FAD ratios and increased lipid desaturation compared to those in the dark zone (Fig. 2n-q). Although these metabolic features were not necessarily co-expressed within the same individual B cells because of the metabolic heterogeneity, their regional enrichment highlights spatially distinct metabolic programs within the GC microenvironment even within the same cell type. These findings corroborate the CODEX cluster analysis (Fig. 2g) and support the notion that B cells in the light zone exhibit a more reduced redox state and anabolic lipid metabolism associated with B cell selection and T cell help, whereas dark zone B cells display lower NADH/FAD and unsaturated lipid ratios, consistent with a more oxidized redox state characteristic of high proliferation and somatic hypermutation. These findings also demonstrated the effectiveness and accuracy of our REDCAT pipeline in mapping single-cell metabolic activity and tissue-level metabolic heterogeneity.

### REDCAT reveals metabolic reprogramming in human lymphoma

Cancer cells that undergo rapid proliferation or progression need to change their metabolic state to meet the increasing demand for energy and biosynthesis ^8,9^. This metabolic reprogramming depletes critical nutrients and generates a hypoxic, acidic tumor microenvironment, which together impair the activation and recruitment of anti-tumor effector immune cells ^62^. To further demonstrate the utility of REDCAT in assessing differential metabolic activities of cancer and immune cells within the tumor microenvironment, we applied this approach to a human lymphoma tissue sample, aiming to uncover novel cell-type–specific metabolic processes that could inform therapeutic strategies targeting metabolic reprogramming. This lymphoma sample includes a region of chronic lymphocytic leukemia (CLL), a low-grade indolent lymphoma, adjacent to a region of diffuse large B-cell lymphoma (DLBCL), a much more clinically aggressive subtype. This clinical phenomenon, rarely captured on the same biopsy, is known as Richter’s transformation (Extended Fig. 3a) ^63^. The zoomed-in H&E image (1.8 mm × 2.0 mm) revealed small, monomorphic tumor cells in the lower region characterized by dense cellular packing, which gradually transitioned into a more loosely packed area containing pleomorphic large tumor lymphocytes in the upper region of the lymphoma sample (Extended Fig. 3a).

To study the metabolic activities in this lymphoma sample, we sequentially acquired NADH, FAD, protein, and lipid signals and generated quantitative radiometric images using TPEF-SRS imaging (Fig 3a, Extended Fig. 3b), followed by CODEX imaging for 35 protein markers in the same ROI (Extended Fig. 4a). Notably, this lymphoma sample exhibited relatively high lipid content observed as scattered puncta with elevated lipid signal intensity (Fig. 3a). Unsupervised clustering of the CODEX data revealed 12 distinct cell clusters, which were then remapped to their original spatial locations to visualize cellular identities within the tissue context (Fig. 3b, c). DLBCL cells, characterized by high expression levels of CD20, PD-L1, and the proliferation marker Ki67, were primarily localized in cluster 11 (Fig. 3b, c). Consistent with the tissue architecture observed in the SRS-protein (CH_3_ vibrational modes) channel, these cells were physically located in the upper region of the ROI (Fig. 3a, Extended Fig. 3b). Morphologically, DLBCL cells displayed large cell bodies with prominent, irregularly shaped nuclei containing small, dense, spherical nucleoli, distinguishing them from other immune cells (Fig. 3d). These features were further validated by quantifying cell size (Extended Fig. 3c, d) and the immunofluorescent intensity of CD20, PD-L1, and Ki67 after cell segmentation (Extended Fig. 3c, e-g), again confirming the histopathological diagnosis. We analyzed the NADH/FAD, lipid/protein, and unsaturated/saturated lipid ratios across CODEX-defined clusters and found that clusters 0 and 1 (B cells, myeloid cells/macrophages and some of the T cells) exhibited significantly high values for lipid and redox ratios (Extended Fig. 3h). These features indicate elevated lipid biosynthetic activity within these clusters. In contrast, the other clusters exhibited no consistent trends or correlations across these metabolic parameters (Extended Fig. 3h), in stark contrast to the coordinated metabolic profiles seen in normal lymph nodes (Fig. 2g). This underscores a pronounced disruption of metabolic homeostasis within the tumor microenvironment. To further investigate single-cell metabolic activity, we performed single-cell annotation (Extended Fig. 3i) and spatially located each individual cell in tissue (Fig. 3e). To quantify the metabolic features, we plotted the histogram distributions of single-cell metabolic ratios (Extended Fig. 5a-c). Notably, the distribution of unsaturated lipid ratios in lymphoma was broader than that in normal lymph nodes (Extended Fig. 5c, 1f), indicating greater unsaturation heterogeneity in lymphoma tissue. This variability may reflect the presence of tumor cells at different stages and their interactions with surrounding cells, which can influence local metabolic states. Consistent with observations in normal lymph nodes, cells in the lymphoma tissue with top-quartile metabolic ratio values (≥0.8 quartile) were predominantly B and T lymphocytes. (Extended Fig. 5d-f). However, unlike normal tissue, tumor cells also make up a substantial fraction of the cells with top-quartile metabolic ratios (Extended Fig. 5e–f), indicating widespread metabolic reprogramming within the tumor niche. When single-cell metabolic features were mapped to their spatial coordinates, cells with high metabolic ratios were broadly dispersed across the tissue but predominantly concentrated in the lower region of the ROI, where CLL cells are the main population (Fig. 3f-h). We noticed metabolic features in the lymphoma tissue lacked the spatial correlation with tissue architecture that was evident in the normal lymph node (Fig. 3f-h, 2o-q). This phenotype may partially be attributed to the disordered “diffuse” structure of lymphoma tissues.

**Fig. 3.**
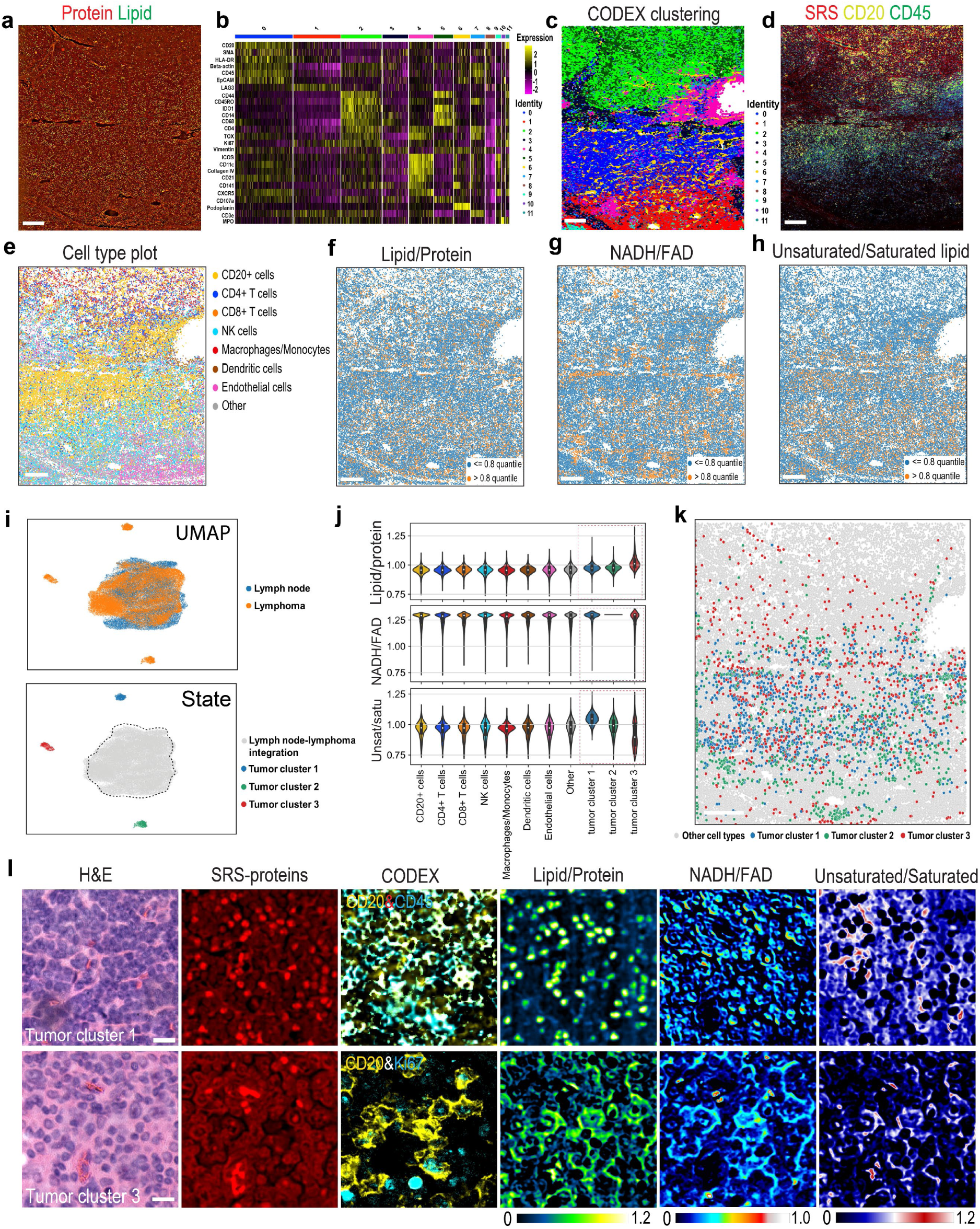
Spatial metabolic and phenotypic profiling of lymphoma using REDCAT. **(a)** SRS image showing protein (red) and lipid (green) distribution in a lymphoma tissue section. Scale bar, 200 μm. **(b)** Heatmap of 35-marker expression profiles across 12 distinct cell clusters identified by CODEX imaging, with cluster identities indicated by the color bar. **(c)** Spatial representation of CODEX unsupervised clustering results in lymphoma tissue. Scale bar, 200 μm. **(d)** Overlay of SRS metabolic imaging with CODEX immunofluorescence for CD20 (yellow), CD3 (cyan), and CD68 (red), showing spatial relationships between metabolic features and immune markers in lymphoma tissue. Scale bar, 200 μm. **(e–h)** Co-registered spatial maps of immune cell types (matrix plot, e), lipid/protein ratio (f), NADH/FAD ratio (g), and unsaturated/saturated lipid ratio (h), highlighting metabolic heterogeneity across cell populations in lymphoma tissue. Scale bars, 200 μm. **(i)** UMAP integrating cell annotations from lymphoma and normal lymph node identified three distinct tumor cell clusters distinguished by their protein expression profiles (top). These three distinct tumor cell clusters have different metabolic states (bottom). **(j)** Violin plots comparing lipid/protein ratio, NADH/FAD ratio, and unsaturated/saturated lipid ratio among different cell types and tumor clusters in lymphoma tissue. **(k)** Spatial distribution of cells assigned to tumor cluster 1 (blue), tumor cluster 2 (green), tumor cluster 3 (red) and other cell types (grey). **(l)** Representative histological and multimodal imaging features of tumor cluster 1 (top) and tumor cluster 3 (bottom), including H&E staining, SRS protein imaging, CODEX immunofluorescence (CD45 or Ki67& CD20), lipid/protein ratio, NADH/FAD ratio, and unsaturated/saturated lipid ratio maps. Scale bars, 20 μm.

To compare immune cell organizations between normal and tumor tissue, we applied spatial neighborhood analysis to both the normal lymph node and the lymphoma sample. In the normal lymph node, spatial neighborhoods exhibited well-defined compartmentalization, corresponding to established anatomical zones (Extended Fig. 5g, h). For example, neighborhood clusters enriched in naïve B cells, plasma cells, and follicular dendritic cells (clusters 2, 3, and 5) were confined to B cell follicles, while macrophage- and dendritic cell–enriched clusters (clusters 0 and 7) were primarily located in the interfollicular and paracortical T cell zones. This spatial organization is consistent with coordinated immune architecture and function in secondary lymphoid organs. By contrast, the B-cell lymphoma sample displayed a disrupted spatial architecture, with a more intermixed and less compartmentalized distribution of spatial neighborhoods (Extended Fig. 5g, h). While some clusters resembling B cell–rich neighborhoods were still present, their boundaries were less distinct. Notably, there was an expansion of macrophage- and NK/T cell–rich neighborhoods (cluster 0 and 1) and a reduction in naïve B cell–dominated clusters, reflecting a shift in tissue composition and immune landscape. This difference was quantitatively captured in the accompanying heatmap (Extended Fig. 5h), which showed altered cell-type enrichments across neighborhoods between normal and tumor tissues. These findings are consistent with the DLBCL being of the activated B-cell subtype, originating from post-germinal center B cells. Lymphoma-associated neighborhoods exhibited increased heterogeneity, with partial replacement of normal immune niches by tumor-associated microenvironments. Together, these data highlight a profound remodeling of immune cell spatial organization in lymphoma, characterized by disrupted follicular structures, altered neighborhood composition, and potential implications for impaired immune metabolism and function.

By integrating metabolic features and protein expression profiles from lymphoma and normal lymph node (Methods), UMAP analysis identified three distinct tumor cell clusters distinguished by their protein expression profiles (Fig. 3i, j). We observed, compared to normal lymph nodes, the lymphoma tissue exhibited globally elevated levels of total lipids, unsaturated lipids, and NADH/FAD redox ratios (Extended Fig. 5i), indicating enhanced lipid metabolism and redox activity in the tumor microenvironment. However, in the lymphoma tissue, compared to other major cell types, tumor cluster 1 exhibited a relatively high unsaturated/saturated lipid ratio, while tumor cluster 3 showed a lower unsaturated lipid ratio but a higher lipid to protein ratio (Fig. 3i). Notably, tumor cluster 3 displayed a heterogeneous distribution of the unsaturated lipid ratio, with a clear bimodal pattern observed in the bar plot (Fig. 3i). This was also confirmed by the UMAP plot (Extended Fig. 5i). Additionally, tumor cluster 2 appears to represent a transitional state between clusters 1 and 3, as its metabolic values lie intermediate between the two (Fig. 3i), potentially reflecting a dynamic shift during the transformation from CLL to DLBCL. These findings suggest that there is a high degree of metabolic heterogeneity between tumor subclusters. To spatially visualize the heterogeneity of metabolic activities in tumor cell clusters, we reprojected the single-cell data onto tissue coordinates and found that tumor cluster 1 predominantly corresponded to CLL cells, whereas tumor cluster 3 was mainly composed of DLBCL cells (Fig. 3k). Zoomed-in images revealed that CLL cells exhibited prominent lipid accumulation (Fig. 3l). These excess lipids, which can be acquired from the microenvironment or synthesized *de novo*, are stored as large lipid clusters. In contrast, DLBCL cells displayed a more diffuse lipid signal in the cytoplasm (Fig. 3l), suggesting an adaptation of lipid metabolism to support rapid membrane biosynthesis and energy demands in the large B cells. Furthermore, we observed a marked elevation in the NADH/FAD redox ratio in DLBCL cells compared to the surrounding non-malignant cells (Fig. 3l), consistent with previous studies showing that DLBCL cells exhibit increased glucose uptake and enhanced glycolytic activity^64^. The dynamic change of metabolic activities between these two tumor subtypes indicates that malignant B cells and their surroundings undergo metabolic reprogramming during transformation from indolent to aggressive lymphoma in order to adapt to the harsh conditions of the tumor microenvironment, such as hypoxia and nutrient deprivation.

Collectively, REDCAT empowers us to spatially elucidate functionally distinct metabolic states whin lymphoma tissues and suggests a possible subpopulation of transitional metabolic state implicated in the transformation from CLL to DLBCL, enhancing our understanding of the metabolic reprogramming in heterogenous cell types within the tumor microenvironment, particularly with regard to tumor evolution.

### REDCAT resolved the change of cell-type–specific chemical profiles during lymphoma transformation

Hyperspectral SRS imaging (HSI) offers the unique advantage to capture subcellular metabolic changes. In HSI stack, each pixel in the image contains a full Raman spectrum, enabling comprehensive chemical profiling at submicron resolution. To fully leverage this capability, we applied 35-panel CODEX and high-resolution SRS-HSI (0.25 µm per pixel, 10 cm⁻¹ spectral resolution) to both normal lymph node and another lymphoma ROI to dissect cell-type**–**specific metabolic profiles during the transformation from low-grade CLL to high-grade DLBCL (Fig. 4a-e, Extended Fig. 6a). Histological examination (H&E staining) confirmed a sharply demarcated boundary between the low-grade and high-grade components within this lymphoma ROI, a rare pathological feature that allowed for precise spatial analysis of metabolic changes across the transformation interface (Extended Fig. 7a). Using unsupervised K-means clustering of pixel-wise SRS spectra, we decomposed the hyperspectral data into 8 distinct chemical clusters (Fig. 4b, c, e). In contrast to normal tissue, the tumor tissue showed enrichment of clusters 4, 6, and 8, characterized by prominent lipid-associated peaks (2850 cm⁻¹ and 2880 cm⁻¹), consistent with increased lipid accumulation shown in Fig. 3a. This observation was also corroborated by single-channel SRS image at 2850 cm⁻¹ (CH₂ symmetric stretch) and lipid radiometric images (Extended Fig. 7b), which showed enriched lipid signals. Notably, SRS-HSI clusters were spatially correlated to the CODEX clusters at single cell level (Fig. 4d, g). To resolve cell-type**–**specific metabolic profile, we integrated HSI results with cell annotations derived from CODEX assay (Extended Fig. 7c, d). We first visualized the spectral cluster enrichment in each cell type by matrix plot and found each cell type contains a wide variety of spectra (Fig. 4h), underscoring the metabolic heterogeneity in tumor tissues. Furthermore, clusters 4, 6, and 8, which were characterized by lipid-associated spectra (Fig. 4c), were enriched across diverse cell types, indicating that lipid remodeling is a widespread feature within the tumor niche. This is consistent with the cell-type**–**specific SRS spectra which showed substantial spectral remodeling in tumor-infiltrating immune cells compared to their normal counterparts (Fig. 4i, j). Consistent with the previous data (Fig. 3g), integration of CODEX-based protein expression with the redox ratio, lipid-to-protein ratio, and unsaturated-to-saturated lipid ratio revealed three tumor clusters in addition to the major cell types shown in the UMAP (Fig. 4k). However, the cell type–oriented clustering pattern was disrupted after embedding the hyperspectral data, further underscoring the metabolic heterogeneity within nominally similar immune cell types (Fig. 4l). Interestingly, three clusters positioned away from the main body emerged after hyperspectral data embedding, suggesting the formation of new cell networks (Fig. 4l).

**Fig. 4.**
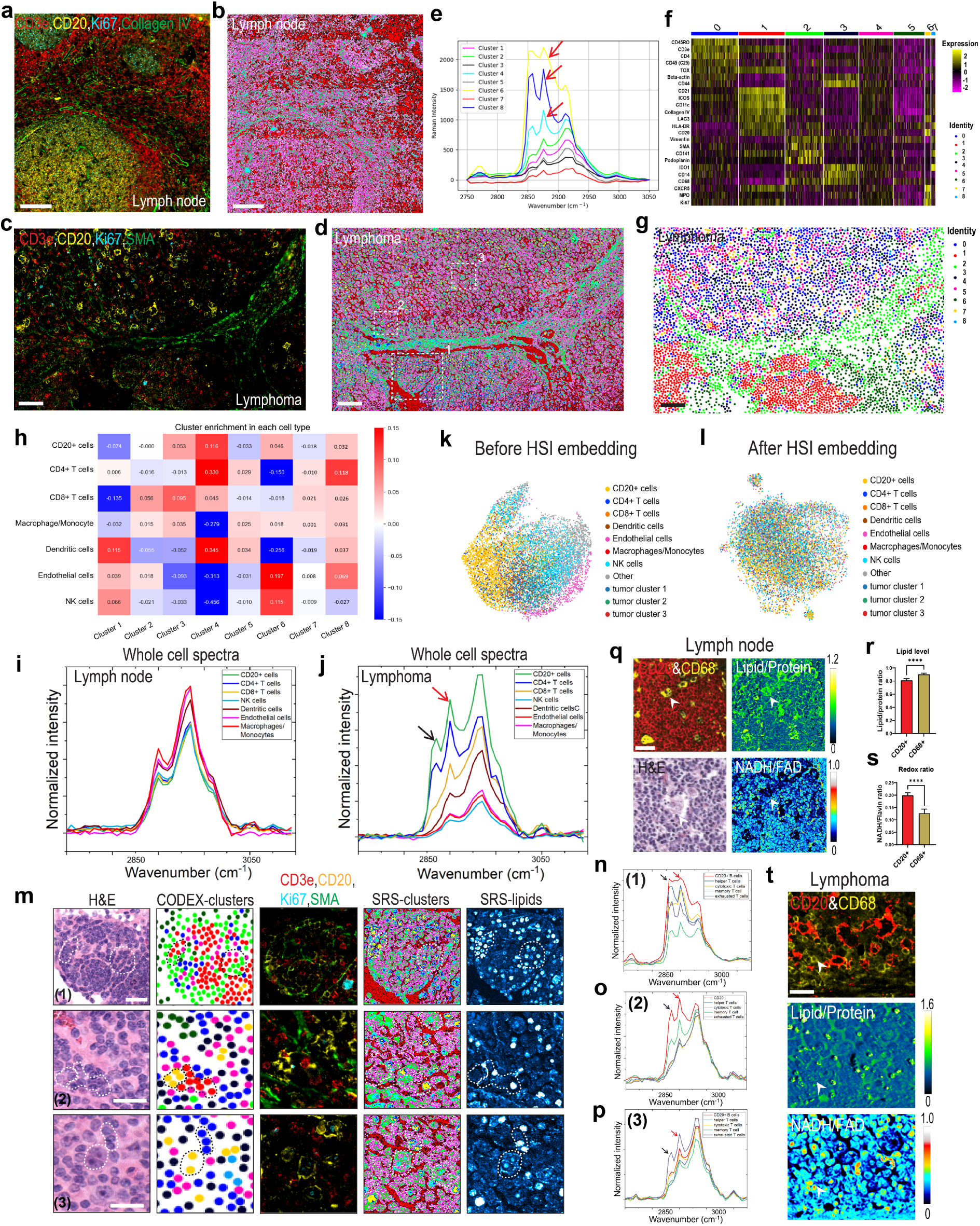
Comparison of metabolic and phenotypic states in normal lymph node and lymphoma. **(a)** CODEX image of human lymph node tissue showing CD3e (red), CD20 (yellow), Ki67 (cyan), and αSMA (red), delineating the major cell types. Scale bar, 100 μm. **(b)** SRS-based spatial k-means clustering map of lymph node tissue. Scale bar, 100 μm. **(c)** CODEX image of a tissue section from transformation of Chronic Lymphoblastic Leukemia (CLL) to Diffuse Large B-Cell Lymphoma (DLBCL), stained for CD3e (red), CD20 (yellow), Ki67 (cyan), and αSMA (green). Scale bar, 100 μm. **(d)** SRS k-means clustering map of the lymphoma sample, with highlighted regions corresponding to CLL (1), intermediate lymphoma (2), and DLBCL (3). Scale bar, 100 μm. **(e)** Integrated mean SRS spectra of k-means–defined clusters in normal lymph node and lymphoma. Red arrows indicate spectra from clusters 4, 6, and 8, which were enriched across multiple cell types in lymphoma. SRS spectra were processed by baseline subtraction and smoothing, followed by normalization to the highest-intensity peaks across the ROI. **(f)** Heatmap of marker expression across clusters in lymphoma, with identity assignments shown in the right color bar. **(g)** CODEX spatial clustering map of lymphoma. Scale bar, 100 μm. **(h)** Heatmap showing enrichment of each immune cell type within k-means spectral clusters in lymphoma. **(i, j)** Average SRS spectra of annotated immune cell types in lymph node (i) and lymphoma (j). In (j) the red arrow indicates the peak at 2880 cm^−1^ which the black arrow points to the peak at 2850 cm^−1^. SRS spectra were processed by baseline subtraction and smoothing, followed by normalization to the highest-intensity peaks across the ROI. **(k, l)** UMAP plots of single-cell populations in lymphoma colored by cell type or tumor cluster before (k) and after (l) HSI embedding. **(m)** Representative multimodal images of selected regions showing H&E, CODEX clusters, CODEX immunofluorescence (CD3e, CD20, Ki67, αSMA), SRS protein channel, and SRS lipid channel (Cyan hot; the brighter the color, the greater the lipid enrichment.). These are the zoomed in views of the regions highlighted in (d), corresponding to (1) CLL region, (2) intermediate region, and (3) DLBCL region. are marked to Lipid accumulation is highlighted in these regions for tumor cells and their surrounding microenvironment. Scale bars, 20 μm. **(n–p)** SRS spectra of cell types from indicated lymphoma regions (1), (2), (3) shown in (m). The red arrow indicates the peak at 2880 cm^−1^ which the black arrow points to the peak at 2850 cm^−1^. SRS spectra were processed by baseline subtraction and smoothing, followed by normalization to the highest-intensity peaks across the respective ROI. **(q)** Representative images in the normal lymph node tissue showing CD20 and CD68 immunostaining (top left), lipid/protein SRS ratio map (top right), H&E staining (bottom left, to show a typical macrophage containing lysosomes), and NADH/FAD redox map (bottom right). The white arrowhead indicates a CD68⁺ macrophage. Scale bar, 20 μm. **(r, s)** Quantification of lipid/protein ratio (r) and NADH/FAD ratio (s) in CD68⁺ macrophages and CD20⁺ B cells in normal lymph node. Values are mean ± SEM. ****p < 0.0001 (Student’s *t*-test). **(t)** Representative images of the DLBCL region showing CD68, CD20 immunostaining (top), lipid/protein SRS ratio map (middle), and NADH/FAD redox map (bottom). The white arrowhead indicates a CD68⁺ macrophage. Scale bar, 20 μm.

To investigate the cell-cell interaction and its function in tumor microenvironment and to postulate whether we can capture the metabolic trajectory of lymphoma transformation from CLL to DLBCL, we arbitrarily defined the CD 20^+^ B cells located between CLL and DLBCL regions as intermediate cells and compared their metabolic profiles to that of large B and CLL cells inside the CLL and DLBCL region, respectively (thereafter CLL and DLBCL cells) (Fig. 4m). We found that CD 20^+^ intermediate cells show a distinct SRS spectra from CLL and DLBCL cells (Fig. 4m-p), especially the lipid-associated peaks at 2850 cm⁻¹ and 2880 cm⁻¹. This result further supports the notion that a putative intermediate state (tumor cluster 2) may exist during CLL to DLBCL transition (Fig. 3i.j). Taken together, these observations support a stepwise metabolic reprogramming model, further refining the previously reported shift from glycolysis to oxidative phosphorylation associated with Richter transformation (ref).

Intriguingly, T cells residing in the three distinct regions also exhibited divergent chemical profiles (Fig. 4m-p, Extended Fig. 7e). These included CD4⁺CD3e⁺ helper T cells, CD8⁺CD3e⁺ cytotoxic T cells, CD4⁺CD8⁺CD45RO⁺ memory T cells, and CD8⁺CD45RO⁺TOX⁺ exhausted T cells. Notably, all T cell subsets in proximity to CLL cells contained substantial lipid accumulation (Fig. 4m, n). This spatial pattern suggests that tumor-induced metabolic remodeling extends beyond malignant B cells to remodel neighboring normal immune cells. The accumulation of lipids in peritumoral T cells may indicate disrupted lipid homeostasis and potential impairment of their effector functions and proliferative capacity.

Notably, we found that, in normal lymph nodes, CD68⁺ macrophages exhibited higher lipid levels and a lower redox ratio than the surrounding CD20⁺ B cells (Fig. 4q, r, s), consistent with active oxidative metabolism, whereas this metabolic pattern was disrupted in lymphoma (Fig. 4t). Such metabolic alterations may facilitate the phenotypic transition from classically activated M1 macrophages to alternatively activated M2 macrophages within the tumor microenvironment ^61,65–67^.

Together, these results demonstrate that REDCAT enables high-resolution chemical profiling of individual cells within complex tissues, revealing both region-specific and cell-type**–**specific metabolic reprogramming during lymphoma transformation.

### Super-resolved REDCAT illuminates subcellular lipid diversity and nuclear heterogeneity in lymphoma

To further investigate cytoplasmic lipid remodeling in lymphoma, we segmented the cytoplasm and identified the pixels with high lipid content based on the SRS-HSI hyperspectral data and mapped the lipid signal to each cell type (Fig. 5a-c). We found that, in normal tissues, lipids are mainly enriched in macrophages, and the lipid spectra exhibited a peak at 2880 cm⁻¹ representing lipid associated vibrational modes (Fig. 5d, e). By contrast, in lymphoma tissues, substantial lipids were found in CD20+ cells (most CLL and DLBCL cells) and identified as different spectral shapes with prominent peaks at both 2850 cm⁻¹ and 2880 cm⁻¹ (Fig. 5d, e). These results were aligned with whole cell spectral analysis in (Fig. 4e).

**Fig. 5.**
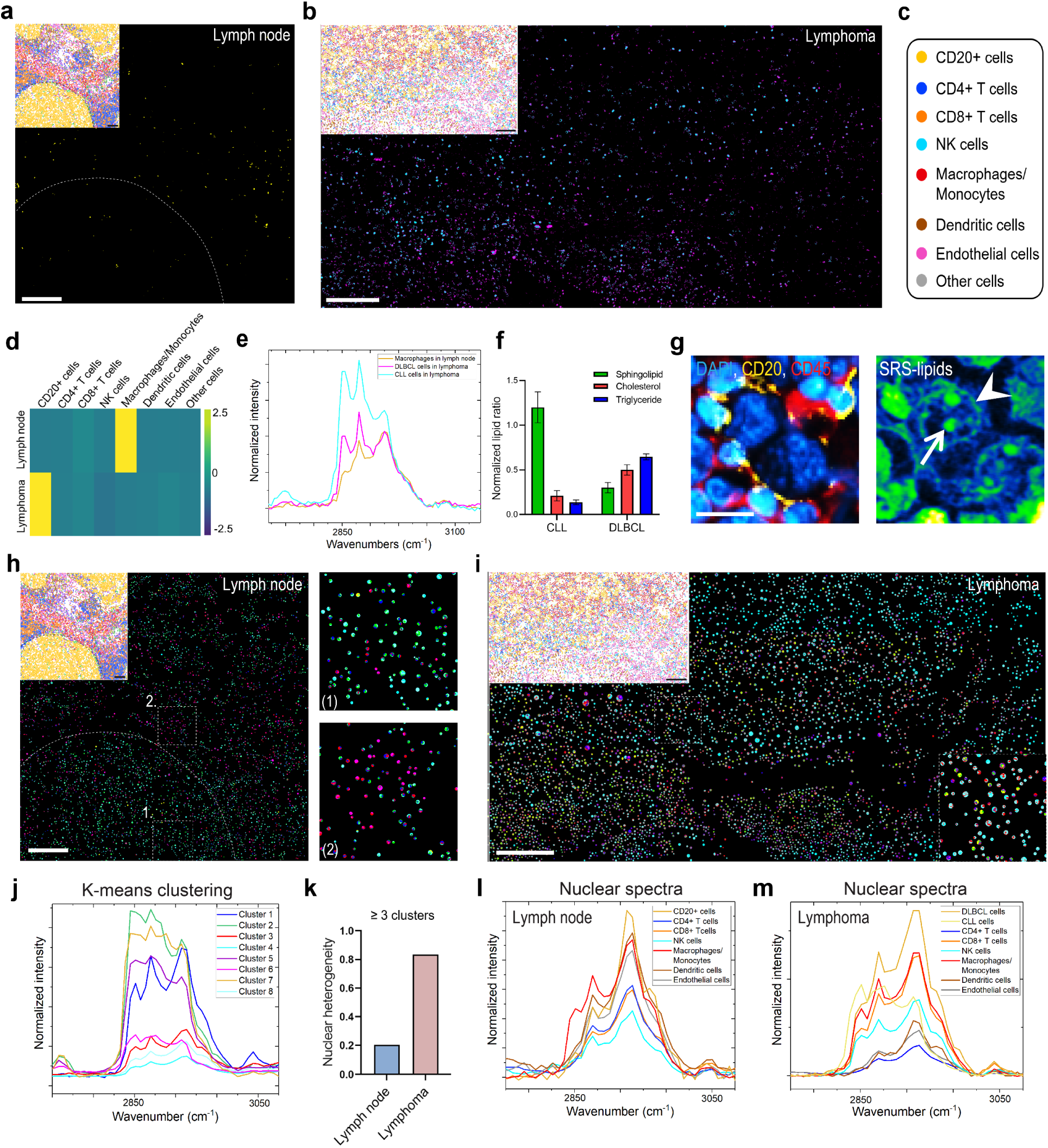
Lipid droplet diversity and nuclear metabolic heterogeneity in normal lymph node and lymphoma. **(a)** Lipid signal in the cytoplasm of cells from normal lymph node tissue showing lipid distribution; inset shows spatial cell type mapping of the same ROI. Scale bars, 100 μm. **(b)** Lipid signal in the cytoplasm of cells from lymphoma tissue showing lipid distribution; inset shows the spatial cell type mapping of the same ROI. Scale bars, 100 μm. **(c)** The different color dots indicate different cell types shown in (a) and (b). **(d)** Heatmap showing the lipid level in each cell type of lymph node and lymphoma. The lipid level was normalized to the total lipids in the corresponding tissue ROI. **(e)** Average SRS lipid spectra of LDs in macrophages in normal lymph node, CLL and DLBCL cells in lymphoma tissue. SRS spectra were processed by baseline subtraction and smoothing, followed by normalization to the highest-intensity peaks of CLL lipid spectra. **(f)** PRM-SRS detection and quantification of the intensity ratios of cholesterol, sphingolipids, and triglycerides to total lipids in CLL and DLBCL cells. **(g)** CODEX image (left) of a region of interest (ROI) showing several large B lymphoma cells and selected markers CD20 (yellow), DAPI (blue), CD45 (red), alongside the SRS image (right) of lipid signal from the same ROI. Arrow indicates nucleolus, arrowhead indicates nuclear envelope. Scale bar, 10 μm. **(h)** Spatial mapping of nuclear SRS spectra in normal lymph tissue; inset shows spatial cell type mapping of the same ROI. Right panels show magnified nuclear clusters inside follicles (1), outside follicles (2). Scale bars, 100 μm. **(i)** Spatial mapping of nuclear SRS spectra in lymphoma; upper inset shows spatial cell type mapping of the same ROI. Lower inset shows the magnified nuclear clusters. Scale bars, 100 μm. **(j)** K-means clusters of nuclear SRS spectra shown in (h) and (i). **(k)** Quantification of the percentage of nuclei displaying ≥3 metabolic clusters in lymph node versus lymphoma. SRS spectra were processed by baseline subtraction and smoothing, followed by normalization to the highest-intensity peaks across the ROI. **(l, m)** Cell-type–specific nuclear SRS spectra of lymph node (l) and lymphoma (m). SRS spectra were processed by baseline subtraction and smoothing, followed by normalization to the highest-intensity peaks across the respective ROI.

A-PoD–assisted SRS super-resolution processing ^68^ illuminated LD structures distinguished by their high density, condensed lipid intensity, and characteristic morphology. It showed CLL cells contained large LD clusters, whereas DLBCL cells harbored significantly smaller LDs (Extended Fig. 8a), suggesting distinct regulation of lipid storage and turnover between tumor subtypes. To investigate the lipid remodeling underlying this transformation, we applied PRM-SRS ^69^ to profile lipid subtypes in CLL and DLBCL cells. The analysis revealed higher levels of sphingolipids in CLL cells, whereas DLBCL cells exhibited elevated levels of cholesterol and triglycerides (Fig 5f, Extended Fig. 8b, c). Intriguingly, super-resolution analysis also uncovered markedly elevated lipid signals in the nucleoli and (peri-)nuclear envelope of DLBCL cells (Fig. 5g and Extended Fig. 8d), suggesting lipids may play a role in gene regulation or the maintenance of nuclear structural integrity. Although both the nucleoli and nuclear envelope exhibited dominant lipid Raman peaks at 2850 cm⁻¹ and 2880 cm⁻¹, their spectral profiles were distinct from each other and more divergent from that of the cytoplasm, reflecting compartment-specific metabolic organization (Extended Fig. 8e).

Motivated by this observation, we next profiled the nuclear metabolic signatures in normal lymph node and lymphoma tissues. Single-nuclear spectrum was extracted and subjected to K-means clustering (Fig. 5h-j). Compared with normal lymph nodes, lymphoma nuclei exhibited greater metabolic heterogeneity, as reflected by a higher number of distinct clusters per nucleus (Fig. 5h, i). Over 80% of lymphoma nuclei contained more than three clusters, whereas only ∼20% of nuclei in normal lymph nodes showed this level of complexity (Fig. 5k). Spatial mapping of the normal lymph node revealed that nuclei within follicles were predominantly enriched in cluster 4, whereas those interfollicular regions were dominated by cluster 6, indicating that nuclear metabolic profiles alone may distinguish functional tissue features (Fig. 5h, j). It indicates nuclear spectra were associated with cell type and subtype. We plotted the average nuclear spectrum for each annotated cell type and observed distinct spectral profiles between cell types in both lymph node and lymphoma. Notably, each cell type–specific spectrum was remodeled in lymphoma compared to lymph node, indicating profound alterations in nuclear metabolism associated with malignant transformation (Fig. 5l, m),

Altogether, REDCAT enables tissue-wise exploration of lipid biology in subcellular compartments including LDs and nuclei. This approach holds promise for detecting subtle lipid metabolic remodeling, identifying diagnostic markers, and tracking the metabolic trajectories of cancer cells during progression from low- to high-grade tumors—offering potential for patient stratification, and personalized treatment.

### REDCAT unveils disorganized collagen architecture and altered composition in lymphoma

In addition to subcellular metabolic profiling, we applied REDCAT to analyze the extracellular matrix by integrating SHG images of collagen fibers (type Ⅰ, Ⅱ, Ⅲ) and CODEX images of collagen fibers (type Ⅳ). As shown in Extended Fig. 9a, in normal lymph nodes, collagen fibers were densely distributed along the capsule and trabeculae, forming a well-organized network. In contrast, lymphoma samples exhibited markedly reduced collagen density and a disorganized fiber network (Extended Fig. 9a).

Morphometric analysis showed that collagen fibers in lymphoma were thinner (Extended Fig. 9b, c) and shorter (Extended Fig. 9d, e) than those in normal lymph nodes. Fiber straightness measurements indicated that lymphoma-associated fibers were more curved, whereas fibers in normal lymph nodes were predominantly straight (Extended Fig. 9f, g). Orientation analysis further revealed a broad angular dispersion of fibers in lymphoma, in contrast to the uniform alignment observed along the capsule and trabeculae in normal lymph nodes (Extended Fig. 9h, i).

SRS spectral profiling of collagen-rich regions uncovered clear biochemical differences between the two tissue types. Lymphoma samples displayed increased peak intensity at 2865 cm⁻¹, corresponding to asymmetric CH₂ stretching features, along with an increased ∼55.6 cm^−1^ of half-height bandwidth (Extended Fig. 9j), indicating changes in collagen composition and molecular packing.

Together, these structural and compositional changes demonstrate that lymphoma is associated with profound remodeling of the extracellular matrix, characterized by collagen fiber loss, disorganization, and altered molecular makeup, which may impact tissue integrity and tumor cell spread and extension into adjacent tissues.

### REDCAT mapping of subcellular metabolic activities in human Liver

To demonstrate the capabilities of the REDCAT platform on to resolved metabolic heterogeneity in the same type of cells as well as compatibility with fresh-frozen (FF) tissues, we applied REDCAT to an FF normal human liver tissue section (Fig. 6, Extended Fig. 10, 11). The liver is organized into hexagonal functional units known as lobules. Nutrient-rich blood flows into each lobule via the hepatic artery and portal vein (PV) at the periportal (PP) region and drains toward the central vein (CV) at the pericentral (PC) region. Based on metabolic functions and capacities, the human liver exhibits functional zonation, with hepatocytes in the periportal zone (zone 1), midzone (zone 2), and pericentral zone (zone 3) differing in their metabolic profiles ^70,71^. Single-cell RNA sequencing has revealed that these gradients drive hepatocytes in different lobule regions to express distinct metabolic genes ^71^, while electron microscopy studies have identified notable differences in mitochondrial morphology between PP and PC hepatocytes ^72^. However, the functional consequences of these structural and molecular differences remain incompletely understood.

**Fig. 6.**
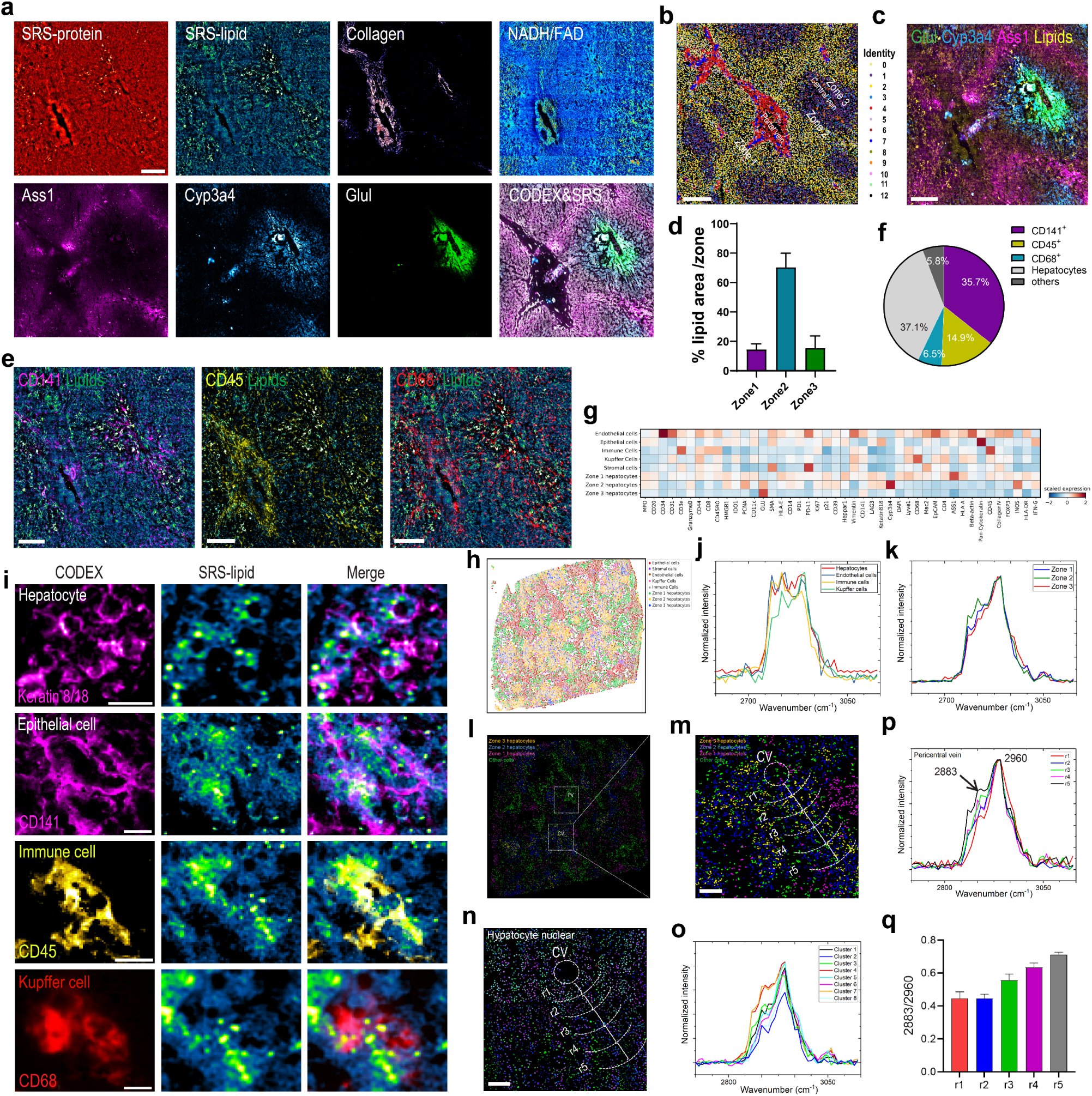
Spatial metabolic and phenotypic mapping of liver zonation using REDCAT. **(a)** REDCAT multimodal imaging of liver tissue showing SRS protein, SRS lipid, SHG collagen, NADH/FAD redox ratio, immunofluorescence for Ass1, Cyp3a4, and Glul, and co-registered CODEX with SRS. Scale bar, 200 μm. **(b)** CODEX clustering map overlaid on tissue architecture highlighting distinct liver zones. Scale bar, 200 μm. **(c)** Spatial distribution of Glul, Cyp3a4, Ass1, and lipid channels mapped onto cell clusters. Scale bar, 200 μm. **(d)** Quantification of lipid area fraction across liver zones 1, 2, 3. **(e)** Spatial co-localization of lipid droplets with CD141, CD45, and CD68 immunostaining. Scale bar, 200 μm. **(f)** Pie chart showing relative abundance of major cell populations (CD141⁺, CD45⁺, CD68⁺, hepatocytes, and others). **(g)** Heatmap of marker expression across annotated cell types in different liver zones. **(h)** Spatial UMAP embedding of single-cell populations colored by cell type. **(i)** Representative high-magnification images of hepatocytes, epithelial cells, immune cells, and Kupffer cells showing cell-type-specific lipid distribution. Scale bars, 20 μm. **(j, k)** Average SRS spectra of annotated cell types in liver zones (j) and from selected immune cell types (k). SRS spectra were processed by baseline subtraction and smoothing, followed by normalization to the highest-intensity peaks across the ROI. **(l, m)** Overview image of the distribution of hepatocytes from different zones highlighting PC shown in (m). Curved lines delineate concentric regions (r1–r5) extending outward from the CV at ∼100 µm intervals, marking nuclei within each zone for spatial classification of hepatocytes. Scale bar, 100 μm. **(n)** Nuclear spatial maps of selected cell populations shown in (m). Scale bar, 100 μm. **(o)** K-means clusters of SRS spectra from nuclei in liver tissue. SRS spectra were processed by baseline subtraction and smoothing, followed by normalization to the highest-intensity peaks across the ROI. **(p)** SRS spectra of indicated regions shown in (n). SRS spectra were processed by baseline subtraction and smoothing, followed by normalization to peak intensity at 2960 cm^−1^. **(q)** Quantification of 2883/2960 cm⁻¹ ratio based on SRS spectra across regions of interest in (n).

To test if these metabolic variations were correlated with spatial positioning of redox ratio and lipid profiles, TPEF-SRS imaging followed by a 47-plexed CODEX imaging was applied to the liver ROI across 1.6 mm × 1.6 mm, containing both central vein and portal vein (Fig. 6a, Extended Fig. 10a, 11a). Unsupervised clustering of CODEX data identified 13 clusters (Extended Fig. 11b), and these clusters were spatially organized to the physiological architecture (Fig. 6b), evidenced by the zonation markers Arginase synthesis 1(ASS1)/zone 1, Cytochrome P450 3A4 (Cyp3A4)/Zone2 and glutamine synthetase (GS or Glul)/Zone3 (Fig. 6a). We detected an uneven distribution of LDs and NADH/FAD ratio among the three zones from TPEF-SRS imaging (Fig. 6a, c, Extended Fig. 11a). This is consistent with literature that the hepatocytes are exposed to different levels of oxidative stress and exhibit different redox states and lipid distribution depending on their relative location within the lobule. Notably, both the LD level and redox ratio were lower in the periportal zone (zone 1) compared with regions adjacent to the central vein (Fig. 6a, c, Extended Fig. 11a). We next quantified the percentage of area occupied by LDs in each zone and found that LDs were predominantly localized to zone 2 (Fig. 6d). By registering LD distribution with cell type–specific markers, we observed that LDs were present not only in ASS1, Cyp3A4, and Glul expressing hepatocytes (37.1%), but also in CD141⁺ (35.7%), CD45⁺ (14.9%), and CD68⁺ (6.5%) cells (Fig. 6c, e, f). Single-cell annotation and spatial mapping (Fig. 6g, h) identified these cells as epithelial, immune, and Kupffer cells, respectively. These lipid-laden cells were mainly situated near the liver sinusoids, forming part of the immune cell landscape of the liver (Extended Fig. 11c). This cell-type**–**specific LD localization was confirmed in another ROI (Extended Fig. 11d).

To investigate cell-type**–**specific LD composition, we acquired SRS-HSI data and conducted chemical profiling on individual LDs. The results revealed that LDs from different cell types exhibited distinct spectral signatures (Fig. 6i, j), suggesting that LD diversity may be critical for defining cell identity and supporting immune function. However, the signaling pathways regulating lipid metabolism within individual cell types, as well as their impact on the liver function, remain to be elucidated. Notably, we also observed differences in the SRS spectral profiles of hepatocytes across liver zonation (Fig. 6k), indicating metabolic heterogeneity associated with spatial location and metabolic state even in the same cell type.

Previous spatial single-nuclear metabolomics studies ^25^ have shown that the PC region is particularly enriched in glucose related metabolites, with concentrations gradually decreasing along the proximal-to-distal axis from the CV. To evaluate whether REDCAT can similarly resolve zonation-dependent nuclear metabolic heterogeneity, we examined the metabolic profiles of individual nuclei within a ROI located near the CV (Fig. 6l, m). SRS-HSI followed by unsupervised clustering identified eight distinct SRS spectral clusters (Fig. 6n, o). Gradient orientation analysis revealed distance-dependent changes in SRS spectra (Fig. 6p), and quantitative analysis further showed that the 2883/2960 ratio increased from the CV toward the outer region (Fig. 6q). However, this phenomenon was not detected near the PP region (Extended Fig. 11e, f), suggesting a region-specific metabolic organization.

Collectively, these findings demonstrate that REDCAT can comprehensively characterize cell-type**–**specific LD diversity and spatial heterogeneity in redox states, lipid distribution, and nuclear metabolic profiles within FF liver tissues.

## Discussion

We developed REDCAT, a multimodal all-optical platform that seamlessly integrates SRS chemical imaging and high-plex CODEX immunofluorescence imaging to enable the spatial exploration of proteomics and metabolic profiles at the single-cell–level in same tissues including clinical archival FFPE samples and FF samples. Application to FFPE lymph node and lymphoma tissues, we demonstrated REDCAT not only tracks intracellular and extracellular metabolic reprogramming during normal immune cell activation but also accurately delineates the subcellular metabolic features during tumor development and progression. Our data indicates that the important lipid species are still preserved in deparaffinized FFPE tissues, likely due to the compact cellular structure that retains small molecules during formalin fixation. This lipid preservation was validated by deparaffinized FFPE 12-month-old tauopathy mouse brain tissues and *Drosophila* brain tissues (data not shown), which still contain substantial lipids. Together, we demonstrated that the capability of REDCAT to spatially explore lipid biology in FFPE tissues, which holds transformative potential for clinical histopathology research.

In normal lymph node tissue, we observed a negative correlation between lipid levels and NADH/ FAD ratios in different cell types. This finding led us to hypothesize that lipid amounts together with redox states could be used to determine the metabolic homeostasis of immune cells in human lymph nodes. However, this correlation is disrupted in lymphoma tissues. In CLL and DLBCL cells, both the global lipid levels, the redox ratio and nuclear heterogeneity are upregulated, indicating significant metabolic remodeling. One possible explanation is that tumor cells are actively engaging the metabolic gene expression and pathway function in catabolism to upregulate NADH coping with the hypoxic stresses while simultaneously *de novo* synthesizing lipid materials for energy source and cellular building blocks. Lipid materials can also result from downregulation of lipogenesis or scavenging from the environment. In CLL, the substantial LD accumulation is consistent with CD36 protein expression reported previously ^73^. Additionally, lysosome abundance is reported to be higher in CLL than normal B cells ^73^, potentially participating in the lipid processing through autophagy. Alternatively, toxic lipids, such as oxidative lipids from membranes or external sources, can be sequestered in LDs, protecting cells from lipotoxicity ^74,75^. However, the metabolic routes of these lipids need further validation. Combining isotope-probed SRS imaging^49^ with genome-wide spatial omics profiling such as Patho-DbiT ^12^ to explore specific lipid metabolism genes in different cell types and transcriptional state can help decipher the source of the lipids and the mechanisms regulating LD generation in normal physiology or tumorigenesis. In fact, we observed that multiple lipid metabolic pathways, along with collagen remodeling associated metabolic pathways, were markedly disturbed within the same lymphoma sample in our DBiTplus study (submitted back-to-back).

We demonstrated unique LD morphology, lipid and nuclear metabolic profiles spatially correlated with CLL and DLBCL cell subpopulations. We observed that the transformation from CLL to DLBCL is accompanied by significant remodeling of lipid distribution, including nuclear and membrane lipids. DLBCL is often associated with chronic inflammation, which can be driven by a persistent inflammatory environment or Epstein-Barr virus (EBV) infection ^63^. In fact, this sample was clinically assessed for EBV and the large, transformed B cells were EBV positive. In sites of chronic inflammation, B cells can acquire genetic changes that promote lipid metabolic reprogramming, which primes tumorigenesis. While this study is exploratory and must be validated by additional focused cohorts, our findings may suggest a potential avenue for targeting abnormal metabolic processes to prevent tumor progression.

CLL and other cell types surrounding the CLL, such as endothelial cells, T cells, NK cells and dendritic cells accumulate more lipids than the distant group in each cell type. This displayed a highly dynamic and heterogenous metabolic remodeling during tumor cell transformation. The result may suggest that cancer cells at different developmental stages engage in extensive crosstalk with the surrounding cells in metabolic modulation and intracellular responses, leading to impaired anti-tumor cell immunity or creating a potentially tumor-promoting immune environment. One possible mechanism is tumor cells release exosomes containing metabolites that are taken up by adjacent cells ^76^. These exosomes can reprogram endothelial cells to produce cytokines that support CLL cell survival and proliferation.

Notably, lipid accumulation was extensively detected within the nucleoli of DLBCL cells. Previous study shows the presence of lipids in nucleoli of mammalian cells ^77,78^. It has been reported that nucleolar protein RRP-8 suppression is linked to abnormal lipid accumulation in nuclei by inducing nucleolar stress ^79^. We hypothesize that nucleolar lipids might be involved in signaling pathways, regulating gene expression, and maintaining the structural integrity of the nucleolus during the large B tumor cell transformation. However, the exact function and mechanisms of their involvement are yet to be further studied.

Beyond the abnormal excess lipid storage in lymphatic cells, other cell types in the tumor environment may exhibit lipodystrophy. In normal macrophages, lipids detected in CD68-positive vesicles are likely derived from debris of endocytosed cells, indicating active macrophage function in maintaining immune surveillance. However, in infiltrated macrophages of lymphoma, lipid levels were reduced, suggesting a dysfunction in processing endocytosed cell debris and a compromised metabolic flux through lysosomes. Investigating why, when, and how lipid metabolism correlates with metabolic features across various cell types in tumor niche is essential for a deeper understanding of tumor pathology in the future.

Similar cell type dependent on LD diversity in size and composition was found in FF healthy liver tissues. Moreover, we found plenty of LDs not just in the parenchymal hepatic cells but in non-parenchymal hepatic cells including immune cells such as Kupffer cells, the resident macrophages involved in immune surveillance. The liver is a major immunological organ that constantly monitors blood from the gastrointestinal tract via the portal vein. Its immune system must maintain tolerance to harmless antigens while rapidly responding to pathogens and injury. LDs in immune cells may be involved in this function. However, how specific lipid profile in each cell type is regulated and how lipid diversity contributes to immune function needs to be further studied. We also found that the nuclear chemical properties showed distance dependent changes along proximal to distal near CV region. It may relate to the genome opening status for the gene expression. To further explore the most related nuclear components or structures representing these spectral changes may help us understand the mechanisms underlying the spatial gene expression as well as the compartmentalized liver function.

Although REDCAT offers a powerful platform for mapping redox, lipid, protein, and nuclear metabolism at single-cell-type resolution in complex biological systems, including clinical tissue specimens, challenges remain in metabolite annotation and coverage. These limitations could be addressed by incorporating hyperspectral acquisition in the fingerprint region, expanding reference libraries for diverse metabolites, and applying generative AI–assisted data analysis. Importantly, these factors do not diminish REDCAT’s current utility. One of REDCAT’s key strengths lies in its ability to identify tissue histological patterns distinguished by high-resolution metabolic features, which can serve as essential clues for subsequent metabolite identification using annotation tool. In addition, REDCAT can be readily integrated with other spatial technologies and is not restricted to any specific MSI platform, broadening its applicability across spatial metabolomics profiling.

In summary, REDCAT offers a high–spatial-resolution, single cell-type–specific metabolic profiling and analysis pipeline that is label-free and requires minimal sample preparation. It is adaptable to diverse biological samples, ranging from cell cultures to complex tissues, and can uncover subtle metabolic alterations that are undetectable or complementary to conventional assays. With future advances in SRS resolution, spectral coverage, and metabolite annotation, REDCAT will enable even more detailed singe-cell or subcellular spatial metabolomic mapping, providing deeper functional and mechanistic insights.

## Materials and methods

### Tissue specimens

De-identified archived human lymphoid tissue samples were obtained from Yale Pathology Tissue Services (YPTS). The tissue retrieval and distribution for research were conducted with the approval of the Yale University Institutional Review Board (Approved IRB #1401013259) and oversight by the Tissue Resource Oversight Committee. Written informed consent for participation in any case where identification was collected alongside the specimen, was obtained from patients or their guardians, in accordance with the principles of the Declaration of Helsinki. Each sample was handled in strict compliance with HIPAA regulations, University Research Policies, Pathology Department diagnostic requirements, and Hospital by-laws.

### Human tissue section preparation

After reviewing all slides by board-certified pathologist and selection of the optimal block, paraffin blocks were sectioned at a thickness of 7-10 μm and mounted on the center of Poly-L-Lysine coated 1 x 3” glass slides. Serial tissue sections were collected simultaneously for TPEF-SRS, CODEX and H&E staining. The sectioning was carried out by YPTS and stored at −80°C until use.

### FFPE tissue information and deparaffinization

The normal lymph node tissue section was acquired from a patient with a BMI value of 32 and the tumor tissue section was collected from a patient with a BMI value of 37. The FFPE and FF tissue sections were processed at Yale University and shipped to UCSD for multimodal SRS imaging workflow. The FFPE tissue slides were bathed with fresh xylene for 5 min and then were transferred to fresh absolute ethanol for 5 min. Followed by incubating 95%, 75%, 50%, and 25% ethanol for 3 min sequentially, the slides were subjected to distilled water for 3 min to be fully hydrated.

### Liver tissue decrosslinking

The FF healthy liver tissue section on the PLL slide followed AKOYA-phynoCycler Fussion FF Tissue Pre-Antibody station. The slide was placed on drierite beads for 5 minutes and placed in Acetone for 10 minutes and let slide sit for 2 minutes at the room temperature. Then the slide was immersed for 2 min in hydration buffer 1 and repeated the same step in hydration buffer 2, finally the tissue on slide was fixed by immersing in 1.6% PFA for 10 minutes. This fixated tissue slide was shipped to UCSD in a falcon tube filled with hydration buffer at 4 °C.

### SRS imaging and high-resolution imaging processing

The SRS images were collected from an upright laser-scanning microscope (DIY multiphoton, Olympus), which was equipped with a 25x water objective (XLPLN, WMP2, 1.05 NA, Olympus). The synchronized pulsed pump beam (tunable 720‒990 nm wavelength, 5–6 ps pulse width, and 80 MHz repetition rate) and stokes beam (wavelength at 1031nm, 6 ps pulse width, and 80 MHz repetition rate) from a picoEmerald system (Applied Physics & Electronics) were coupled and introduced into the microscope. Upon interacting with the samples, the transmitted signal from the pump and Stokes beams was collected by a high NA oil condenser (1.4 NA). A shortpass filter (950 nm, Thorlabs) was used to completely block the Stokes beam and transmit the pump beam only onto a Si photodiode array for detecting the stimulated Raman loss signal. The output current from the photodiode array was terminated, filtered, and demodulated by a lock-in amplifier at 20 MHz. The demodulated signal was fed into the FV3000 software module FV-OSR (Olympus) to form images using laser scanning. SRS protein, lipid, unsaturated lipid and saturated lipid signal were obtained at vibrational modes 2930 cm^−1^, 2850 cm^−1^, 2880 cm^−1^ and 3015 cm^−1^ in a single tile of 512 x 512 pixels, at dwell time 40 μs and 100 tiles were stitched to generate the large tissue mapping. To quantify the unsaturated lipid ratio, the intensity of C=C vibrational modes (peak intensity at 3015 cm^−1^) was divided by the intensity of saturated vibrational modes (peak intensity at 2880 cm^−1^), resulting in ratiometric images. Minor intensity adjustment (brightness and/or contrast) was performed in ImageJ. The figures were assembled in Adobe Illustrator 2023. To reach a high resolution of the subcellular organelles, the measured images can be processed by Adam based Poinitillism Deconvolution (A-PoD). Based on a function to describe blurring pattern of point emitters, so called point spread function, the super-resolution images can be reconstructed.

### SRS hyperspectral imaging and clustering analysis

SRS hyperspectral images were collected from the same setup described above. The hyperspectral imaging stacks were taken with 51 spectral steps at 40 μs pixel dwell time, covering the whole CH stretching region from 2800 to 3150 cm^−1^. The intensity profiles of the hyperspectral stacks from the regions of interest were plotted in ImageJ and further processed by baseline correction, smoothing and normalization in OriginPro 2017. SRS hyperspectral imaging of 50 mM Rhodamine 6G dye (R6G) solution was performed in each imaging session for laser power calibration. The spectrum of R6G was divided by all the sample spectra to obtain intensity-normalized SRS spectra for quantification purposes. Unsupervised K-means clustering method was applied to generate metabolic mapping at each pixel. SRS spectra were normalized to the highest-intensity peaks across the respective ROI a to obtain intensity-normalized SRS spectra shown in all pixel-wise clustering and region/cell-type–specific spectra figures.

### Optical redox imaging

The two-photon fluorescent microscopy was used to collect the NADH and FAD signals from the tissue. The NADH signal was excited by 780 nm, the emission wavelength was collected at 460 nm. The FAD signal was excited by 860 nm, the emission wavelength was collected at 515 nm. The ratiometric images were generated by NADH /FAD. The ratio was further quantified in ImageJ and plotted in GraphPad 10.

### CODEX spatial phenotyping using PhenoCycler-Fusion

After receiving the slides following the SRS workflow, a modified version of the CODEX PhenoCycler-Fusion protocol (Link to protocol) was adopted. Since the tissue had already been deparaffinized and rehydrated during the SRS workflow, the CODEX process began with a gentle antigen retrieval step using 1X AR9 buffer for 5-10 minutes. The tissue was then allowed to cool to room temperature and was rinsed twice with nuclease-free water and hydration buffer, followed by staining buffer as the antibody cocktail was prepared. The tissue slide was incubated with the antibody cocktail at room temperature for 3 hours in a humidity chamber. After incubation, the tissue underwent a series of steps including post-fixation, ice-cold methanol incubation, and a final fixation step.Attached to the flow cell, the tissue section was incubated in 1X PhenoCycler buffer with additive for at least 10 minutes to improve adhesion. The CODEX cycles were then set up, the reporter plate was prepared and loaded, and the imaging process began. A final qptiff file was generated at the end which could be viewed using QuPath V0.5.168. For further details on the PhenoCycler antibody panels, experimental cycle design, and reporter plate volumes, see Table - Codex panel details.

### Flow cell removal and H&E staining

Following the CODEX imaging workflow, the flow cell can be removed, and histological H&E staining performed on the same tissue section. To remove the flow cell, the tissue slide with the flow cell is immersed in xylene or HistoClear for a minimum of 20 minutes to weaken the adhesive. A razor was then used to carefully detach the flow cell from the tissue slide. The tissue slide was then rinsed thoroughly with deionized water and then with 1X PhenoCycler Buffer without additive three times for 10 minutes each. Histological H&E staining on the FFPE sections was conducted by YPTS.

### Preprocessing of CODEX data

Cell segmentation was performed using a StarDist-based model ^80^ in QuPath ^81^. The DAPI channel served as the nuclear marker for cell segmentation. The mean intensity of each marker per segmented cell was exported together with the centroids of each cell as a CSV. Downstream analysis was performed using Seurat 4.3.0 package. The dataset was normalized and scaled using NormalizeData and ScaleData functions. Linear dimensionality reduction was then performed with the “RunPCA” function. The “FindNeighbors” function embedded spots into a K-nearest neighbor graph structure based on Euclidean distance in PCA space, and the “FindClusters” function was used to cluster the spots. The “RunUMAP” function was used to visually show spatial heterogeneities through the Uniform Manifold Approximation and Projection (UMAP) algorithm. The clusters were plotted spatially using the ImageDimPlot function. The FindAllMarkers function was used to find the differentially expressed proteins in each cluster and the heatmap plotted using the DoHeatMap function.

### Quality control and preprocessing for single cell analysis

Cell and nuclei boundaries were generated using the Mesmer ^82^ with pretrained weights. The DAPI channel was employed to identify nuclei, while the CD45 channel marked the cell membranes. The default resolution of 0.5 micrometers per spot was used for cell mask prediction. Only cells with sizes falling within the [0.05, 0.95] quantile range and exhibiting DAPI intensities above the 0.1 quantile threshold were included in the analysis. CODEX feature extraction involved summing the signal for each feature across individual cells. For each feature, the 0.05 and 0.95 quantiles were computed, and the values were normalized to fit within a [0, 1] scale, where 0 corresponded to the 0.05 quantile and 1 to the 0.95 quantile. Values outside this range were capped at 0 or 1, as needed.

### Cell type annotation

Cell type annotation was performed using the Astir package ^83^. Cleaned and normalized protein expression matrices were used as input, ensuring high-quality data for downstream analysis. For main Figures 3, 4, 5, marker genes used in annotation included B cells (CD45, CD20, CD21), CD4+ T cells (CD45, CD3e, CD4), CD8+ T cells (CD45, CD3e, CD8), NK cells (CD45, CD107a), macrophages (CD45, CD68, CD163), monocytes (CD45, CD14), dendritic cells (CD45, CD141), and endothelial cells (CD31). For Main Figure 6, marker genes used in annotation included Zone 1 hepatocytes (ASS1, HLA-A), Zone 2 hepatocytes (Cyp3a4), Zone 3 hepatocytes (GLU), Epithelial cells (EpCAM, Pan-Cytokeratin), Stromal cells (PD-L1), Endothelial cells (Lyve1, CD31, CD34), Kupffer Cells (CD68), Immune Cells (CD45). The annotation process was initialized with the fit_state() function, using default parameters to estimate an initial probabilistic state for each cell. The fit() function was then applied to train the model, which estimates the posterior probability of each cell belonging to predefined cell types based on marker expression patterns. Cells were annotated with the type showing the highest posterior probability. After cell annotation, SRS single cells were mapped onto their matched CODEX cells and the metabolic profiles were extracted for analysis.

### MaxFuse integration of CODEX and scRNA-seq data

For main Figures 2j and 2n, we obtained the human lymph node scRNA-seq dataset from the Cell2location repository (available for download at Link to reference dataset). Before integration, all scRNA-seq data underwent standard preprocessing in ScanPy, including library-size normalization, log1p transformation, and selection of highly variable genes, retaining the 5,000 most variable genes. Integration of the scRNA-seq and CODEX modalities was performed with the MaxFuse algorithm ^54^. Cross-modal feature links were defined by matching protein and gene names; among these, features with standard deviation > 0.01 were kept enhancing integration performance. For pivot matching, the number of principal components used to construct the nearest-neighbor graph was chosen using the elbow criterion on the singular-value decomposition (SVD). To accommodate the weak linkage between modalities, a smoothing weight of 0.3 was applied as recommended by MaxFuse ^54^. After integration, scRNA-seq cells were mapped onto their matched CODEX cells and visualized in spatial plots.

### SRS single cell analysis

Fixed tissue sections were imaged by multimodal SRS microscopy and CODEX sequentially. Because fixation and gentle handling minimize mechanical drift, we assumed no appreciable non-rigid deformation during imaging. Any residual field-of-view shifts, rotations, and uniform scaling were treated as global rigid/affine effects and corrected computationally. To co-register the modalities, we manually selected a set of visually reliable control points in the SRS and CODEX images. Given the two coordinate arrays, we estimated an optimal affine transformation by least squares using skimage.transform.estimate_transform(), which minimizes the sum of squared distances between matched control points. The SRS image was mapped into the CODEX coordinate system by applying the inverse transform with skimage.transform.warp(), yielding pixel-level alignment suitable for downstream cell-based analysis. Mapped cell centroids were then generated accordingly so that per-cell measurements could be compared one-to-one across SRS and CODEX. At the single-cell level, we computed ratio metrics reflecting metabolic and biochemical state, including NADH/FAD (redox balance), Lipid/Protein (overall lipid enrichment), and Unsaturated-to-Saturated lipid (degree of lipid unsaturation). For each metric, we summarized the upper tail by the 0.8 quantile to capture high-value distributions while remaining robust to occasional extremes. Scatter plots were generated to visualize the spatial distribution of the computed ratios across the tissue. In practice, affine alignment is generally sufficient for fixed sections acquired in close succession, providing a transparent, low-variance correction. Subtle local warping at tissue edges or folds may benefit from higher-order models, which could be worth exploring in cases with pronounced distortion.

## ACKNOWLEDGMENTS

We acknowledge the support received from the U.S. National Institutes of Health (NIH) including grants NIH R01GM149976, U01AI167892, 5R01NS111039, R21NS125395, U54DK134301, U54HL165443 (all to L. S.), U54CA274509, U54CA268083, UH3CA257393, RF1MH128876, U54AG079759, U54AG076043, R01CA245313, RM1MH132648 (all to R.F.). and U01CA294514 (to R.F., M.X., and Z.M.). Dr. Lingyan Shi (L.S.) is a Sloan Research Fellow. Z.M. is supported by NSF awards 2345215 and 2245575. We also thank Yale West Campus cleanroom team for assistance with microfluidic wafer fabrications and the YPTS team for FFPE tissue sectioning and staining. Computational data analysis was conducted with the support of Yale High Performance Computing clusters (HPC).

## Author Contributions

Conceptualization: L.S. and R.F.; Methodology: Y. L., Z. Z., A. E., R. F., Z. M. and L. S.; Experimental Investigation: Y. L., A. E., J. N., X. Q. and N. F.; Data Analysis: Y. L., Z. Z., A. E., J. V., A. A. F. H. J. and Z. B.; Data Interpretation: M. L. X., R. F.,L. S. and Z. M.; Resources and valuable inputs: Y. L., A. E., Z. Z., M. L. X., N. F., J. N., X. Q. and N. Z. Original Draft: Y.L., A.E., Z.Z. All authors reviewed, edited, and approved the manuscript.

## Extended Figures

**Extended Figure 1.**
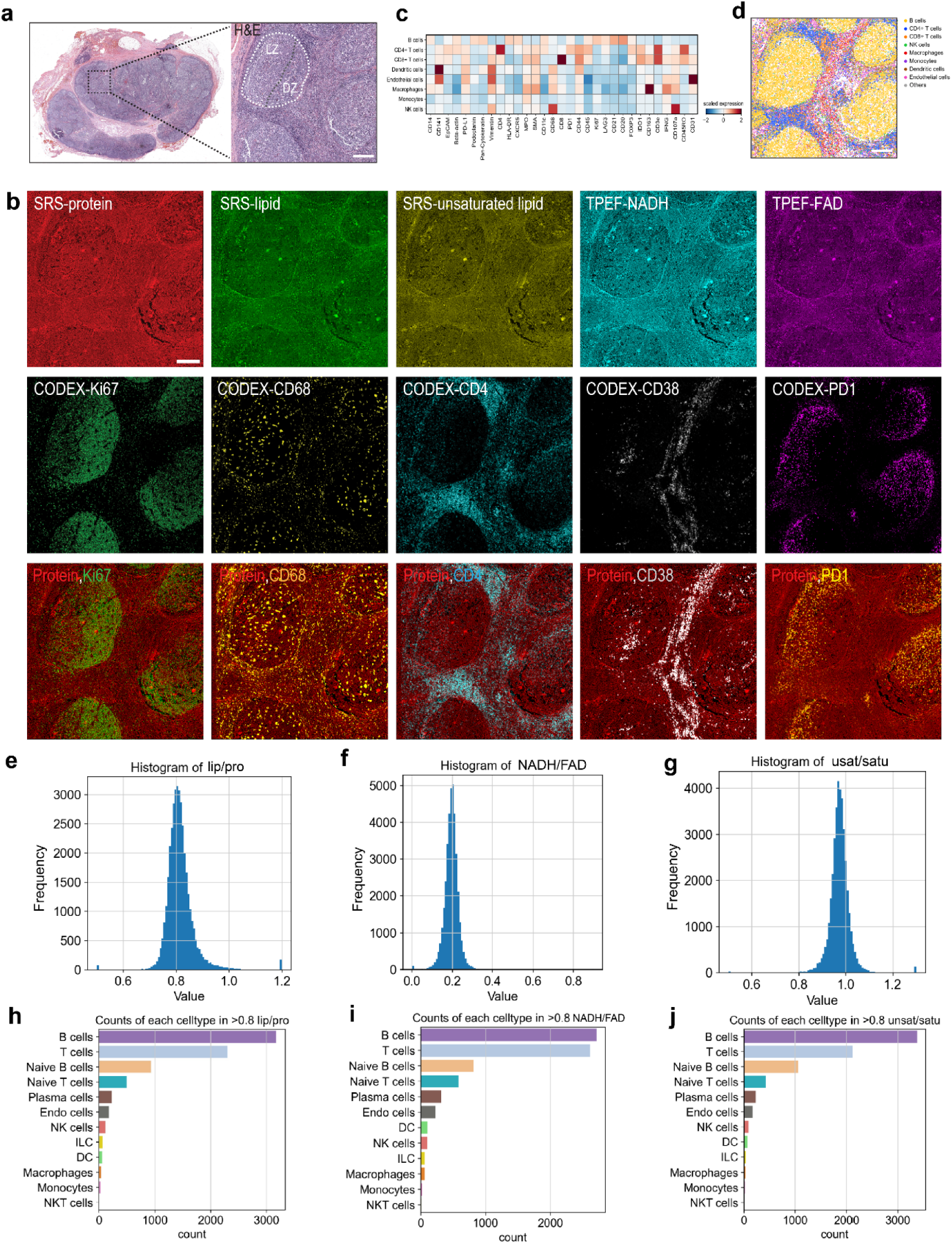
Multimodal metabolic and proteomic imaging of lymphoid tissue using REDCAT. **(a)** H&E-stained whole-section and magnified views showing the tissue architecture of normal lymph node. The geminal center (GC) light zone (LZ) and dark zone (DZ) were highlighted. Scale bar, 200 μm. **(b)** Multichannel imaging of the same tissue section: SRS-protein, SRS-lipid, SRS-unsaturated lipid, TPEF-NADH, and TPE-FAD autofluorescence; CODEX immunofluorescence for Ki67, CD68, CD4, CD38, and PD-1; and corresponding merged overlays of SRS protein with each immunostaining channel. Scale bar, 200 μm. **(c)** Heatmap showing scaled expression of selected protein markers across annotated immune cell types in normal lymph node. **(d)** Spatial mapping of annotated immune cell types based on protein expression in normal lymph node. Scale bar, 200 μm. **(e–g)** Histograms of lipid/protein ratio (e), NADH/FAD ratio (f), and unsaturated/saturated lipid ratio (g) for all single cells in normal lymph node. **(h–j)** Counts of each annotated cell type exceeding thresholds of lipid/protein ratio ≥0.8 quartile (h), NADH/FAD ratio ≥0.8 quartile (i), and unsaturated/saturated lipid ratio ≥0.8 quartile 8 (j), highlighting cell populations with metabolic ratios in the top quantile.

**Extended Figure 2.**
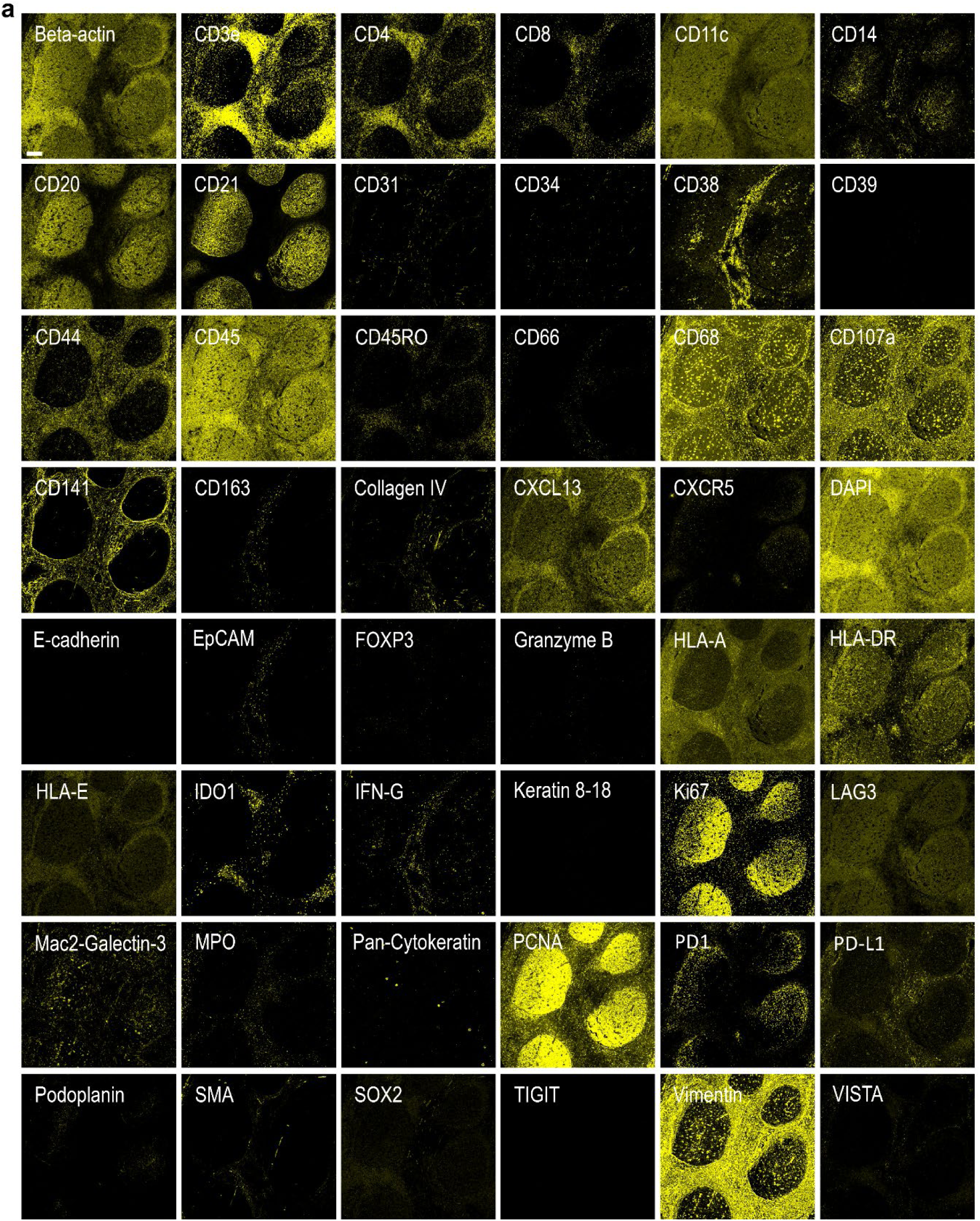
CODEX antibody panel validation in tissue sections. **(a)** Representative single-channel images from the 48-plex CODEX antibody panel showing specific staining patterns for immune, stromal, epithelial, and functional markers. Each panel corresponds to one antibody target (label indicated) acquired from the same tissue section. Markers include structural proteins (e.g., Beta-actin, Vimentin), immune lineage markers (e.g., CD3e, CD4, CD8, CD11c, CD14, CD20, CD21, CD38, CD45, CD68, CXCR5), stromal and extracellular matrix markers (e.g., Collagen IV, SMA, Podoplanin), checkpoint/inhibitory molecules (e.g., PD-1, PD-L1, TIGIT, LAG3, VISTA), and functional molecules (e.g., Ki67, IFN-γ, Granzyme B, IDO1). DAPI channel shows nuclear counterstaining. All images were acquired under identical imaging conditions to ensure comparability of staining patterns. Scale bar, 200 μm.

**Extended Figure 3.**
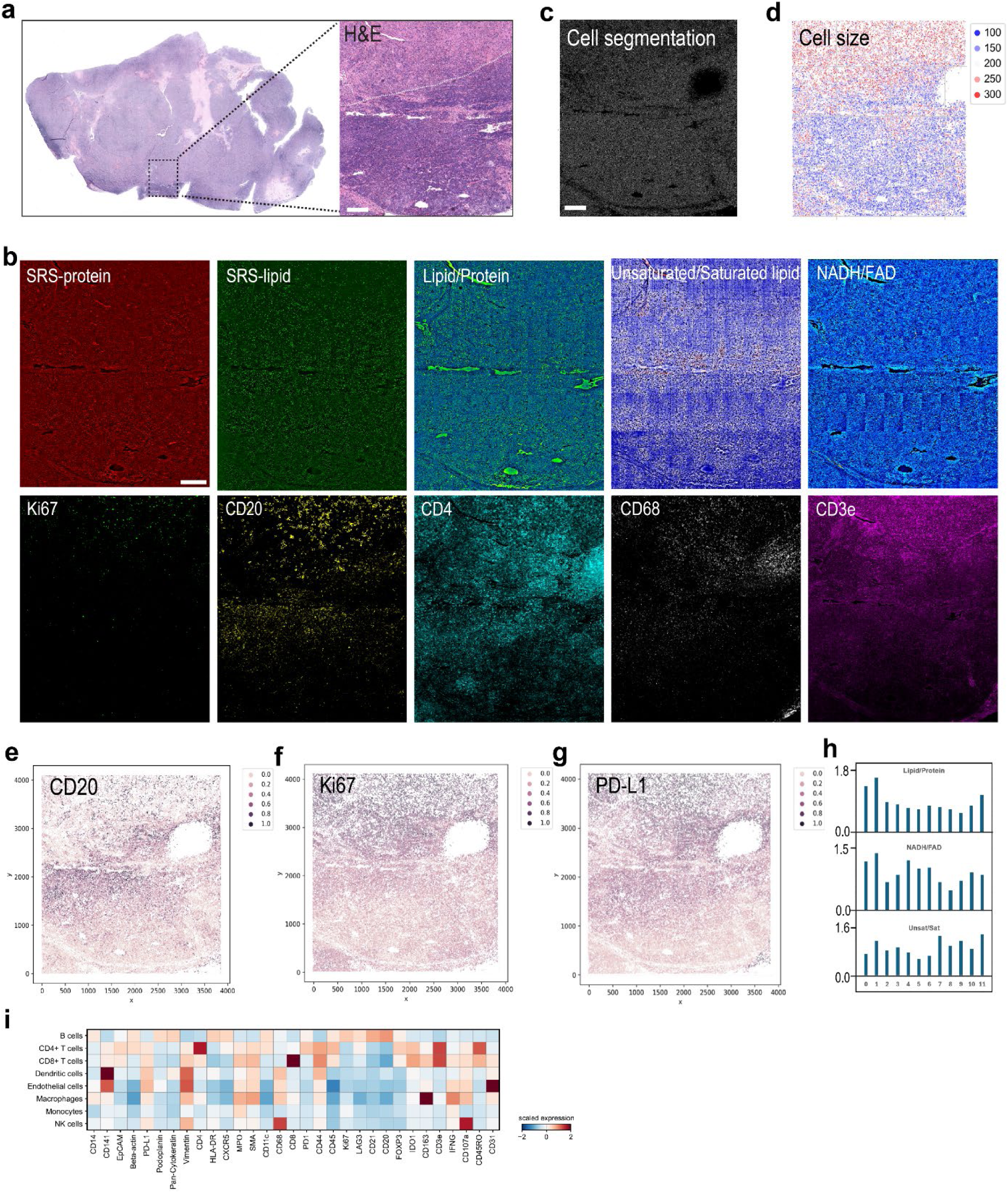
Multimodal metabolic and proteomic imaging of lymphoma tissue. **(a)** Whole-section and magnified H&E images of lymphoma ROI showing tumor architecture. Scale bar, 200 μm. **(b)** Multimodal imaging including SRS-protein, SRS-lipid, lipid/protein ratio, unsaturated/saturated lipid ratio, NADH/FAD ratio, and CODEX immunofluorescence for Ki67, CD20, CD4, CD68, and CD3e. Scale bar, 200 μm. **(c)** Cell segmentation of the same region shown in (a, b) to quantify the cell size. Scale bar, 200 μm. **(d)** Spatial map of cell size distribution across the lymphoma ROI shown in (a, b). **(e–g)** Whole-slide spatial distribution maps of CD20 (e), Ki67 (f), and PD-L1 (g) expression derived from CODEX imaging. **(h)** Quantification of lipid/protein ratio, NADH/FAD ratio, and unsaturated/saturated lipid ratio across CODEX identified cell clusters of the lymphoma ROI. **(i)** Heatmap of marker expression across annotated cell types in the lymphoma ROI.

**Extended Figure 4.**
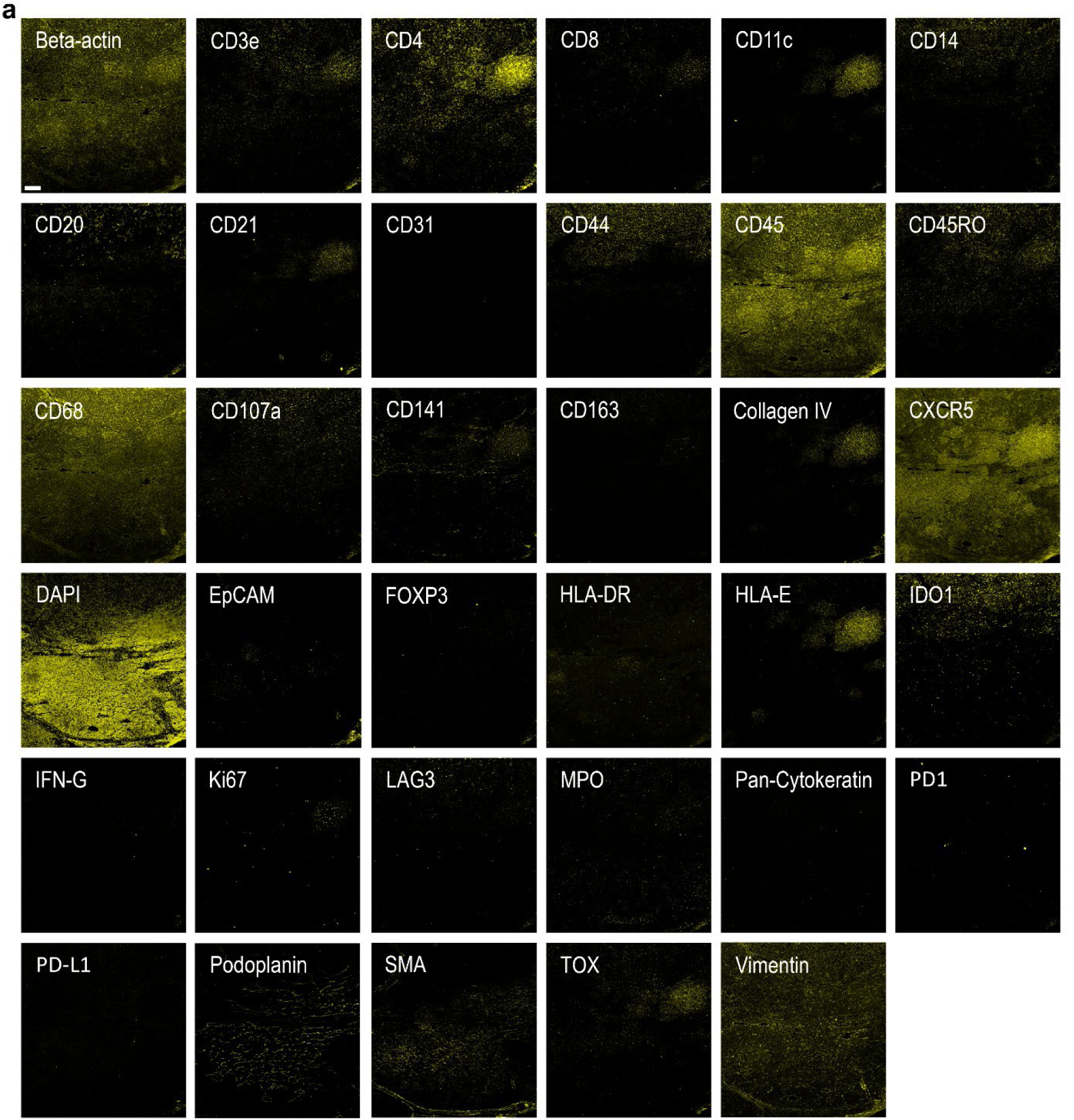
CODEX antibody panel validation in tissue sections. **(a)** Representative single-channel images from the CODEX antibody panel showing staining patterns for structural, immune, stromal, and functional markers. Markers include Beta-actin, immune lineage markers (e.g., CD3e, CD4, CD8, CD11c, CD14, CD20, CD21, CD44, CD45, CD45RO, CD68, CD107a, CD141, CD163, CXCR5), stromal/extracellular matrix proteins (e.g., Collagen IV, SMA, Podoplanin, Vimentin), checkpoint/inhibitory molecules (e.g., PD-1, PD-L1, LAG3, TOX, IDO1), and functional markers (e.g., Ki67, FOXP3, IFN-γ, MPO). The DAPI channel is included for nuclear counterstaining. All images were acquired under identical imaging conditions for direct comparison of staining specificity and intensity. Scale bar, 200 μm.

**Extended Figure 5.**
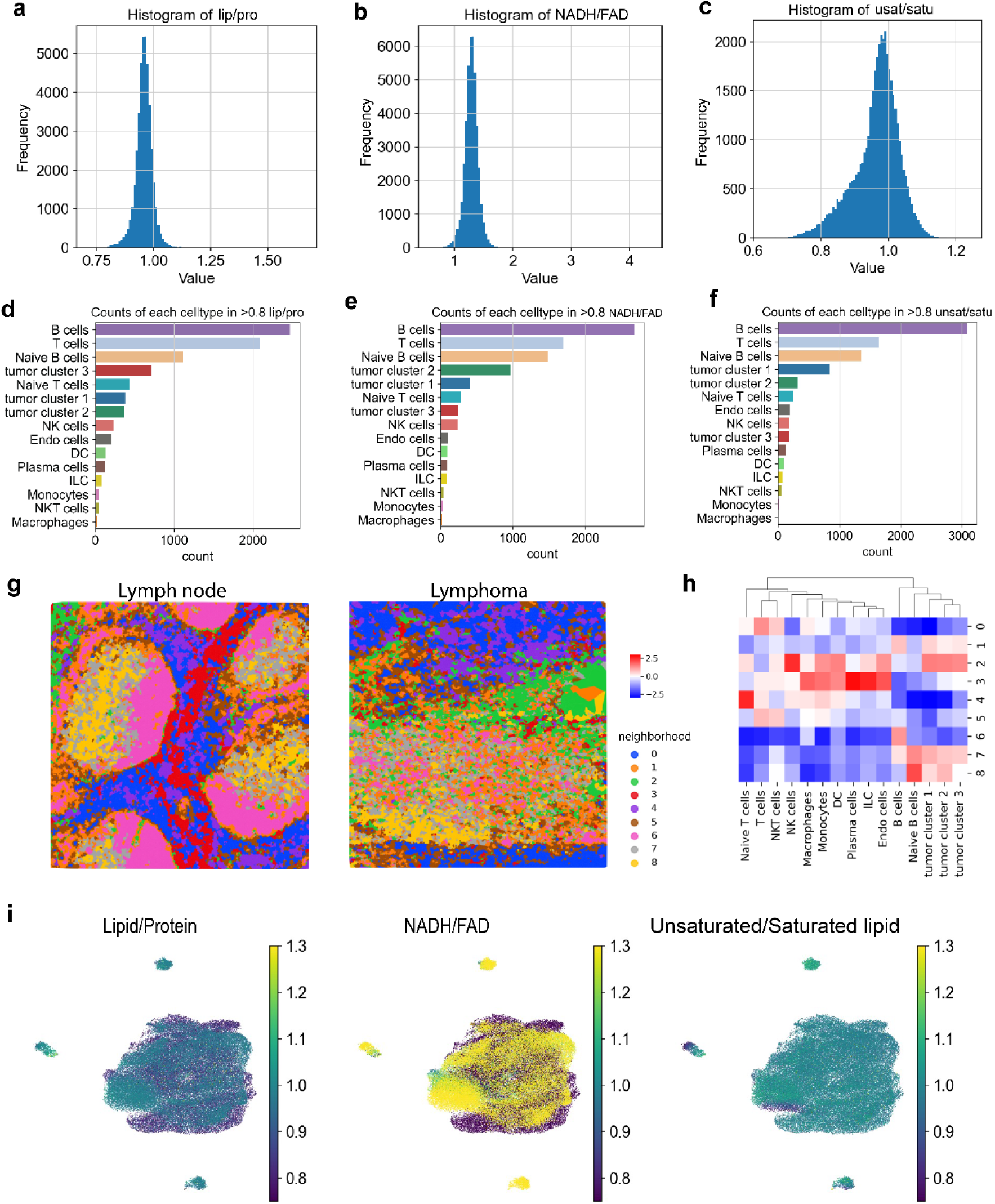
Single-cell metabolic profiling and neighborhood analysis of lymph node and lymphoma. **(a–c)** Histograms of lipid/protein (c), NADH/FAD (d), and unsaturated/saturated lipid (e) ratios for all single cells in lymphoma. **(d–f)** Counts of each annotated cell type with metabolic ratios greater than 0.8 for lipid/protein (f), NADH/FAD (g), and unsaturated/saturated lipid (h), highlighting metabolically distinct subsets in lymphoma**. (g)** Spatial neighborhood maps for lymph node (left) and lymphoma (right), showing cellular neighborhood composition. **(h)** Heatmap of neighborhood enrichment scores across cell types and tumor clusters, identifying microenvironmental differences between lymph node and lymphoma. **(i)** UMAP feature plots showing single-cell lipid/protein ratio, NADH/FAD ratio, and unsaturated/saturated lipid ratio in both lymph node and lymphoma.

**Extended Figure 6.**
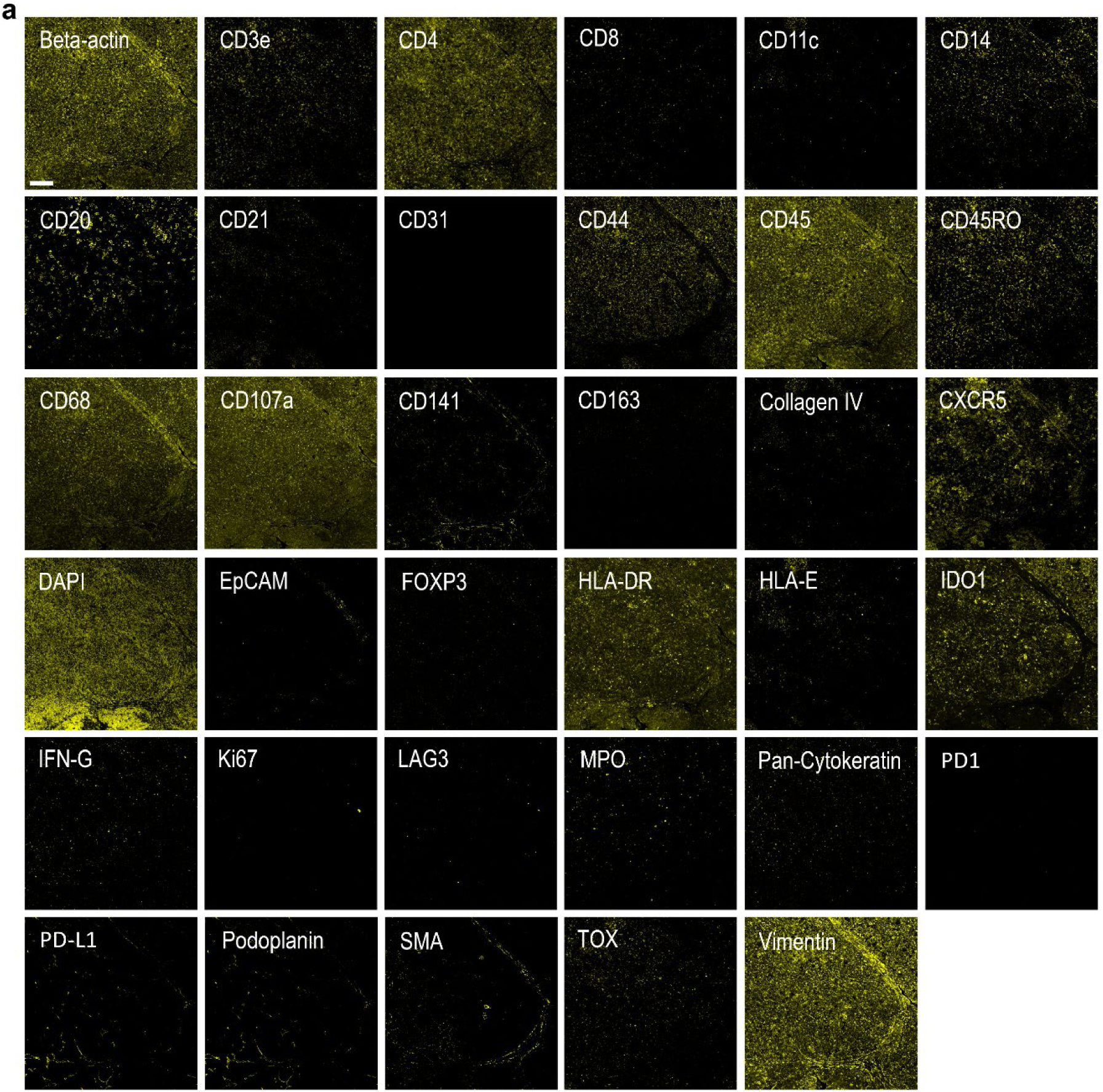
CODEX antibody panel validation in lymphoma tissue. **(a)** Representative single-channel images from the 35-plex CODEX antibody panel showing staining specificity for structural, immune, stromal, and functional markers. Markers include Beta-actin, immune lineage markers (e.g., CD3e, CD4, CD8, CD11c, CD14, CD20, CD21, CD44, CD45, CD45RO, CD68, CD107a, CD141, CD163, CXCR5), stromal/extracellular matrix proteins (e.g., Collagen IV, SMA, Podoplanin, Vimentin), checkpoint/inhibitory molecules (e.g., PD-1, PD-L1, LAG3, TOX, IDO1), and functional/activation markers (e.g., Ki67, FOXP3, IFN-γ, MPO). DAPI staining is included for nuclear visualization. All channels were imaged under identical acquisition conditions to ensure comparability of staining intensity and localization. Scale bar, 200 μm.

**Extended Figure 7.**
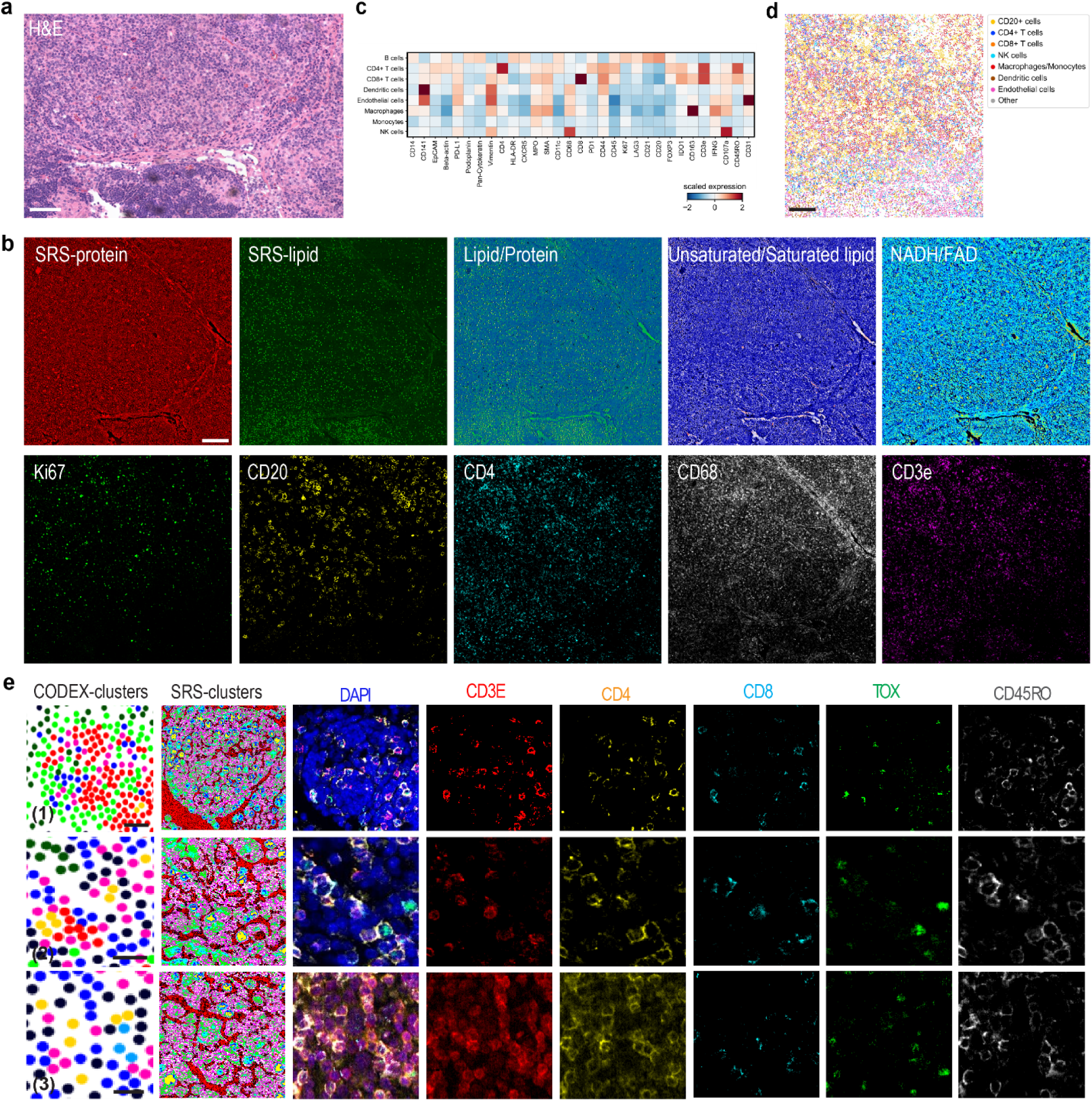
Multimodal metabolic and proteomic characterization of immune populations in lymphoma tissue. **(a)** H&E image of the lymphoma ROI containing low-grade and high-grade components in the lymphoma tissue, as well as a transformation interface between these two compartments. Scale bar, 200 μm. **(b)** Heatmap of scaled expression values for selected markers across annotated immune cell types, including CD4⁺ T cells, CD8⁺ T cells, CD20⁺ B cells, dendritic cells, endothelial cells, macrophages/monocytes, and NK cells in the lymphoma ROI. Scale bar, 200 μm. **(c)** Spatial map of annotated immune cell types across the lymphoma ROI. **(d)** Co-registered SRS and CODEX imaging of the same lymphoma ROI showing SRS-protein, SRS-lipid, lipid/protein ratio, unsaturated/saturated lipid ratio, NADH/FAD ratio, and CODEX immunofluorescence for Ki67, CD20, CD4, CD68, and CD3e. Scale bar, 200 μm. **(e)** Representative regions from three distinct microenvironments (Fig. 4m) showing CODEX cluster assignments, corresponding SRS cluster overlays, and high-magnification immunofluorescence for DAPI, CD3e, CD4, CD8, TOX, and CD45RO, illustrating the major T cell subtypes and spatial organization. Scale bars, 20 μm.

**Extended Fig. 8.**
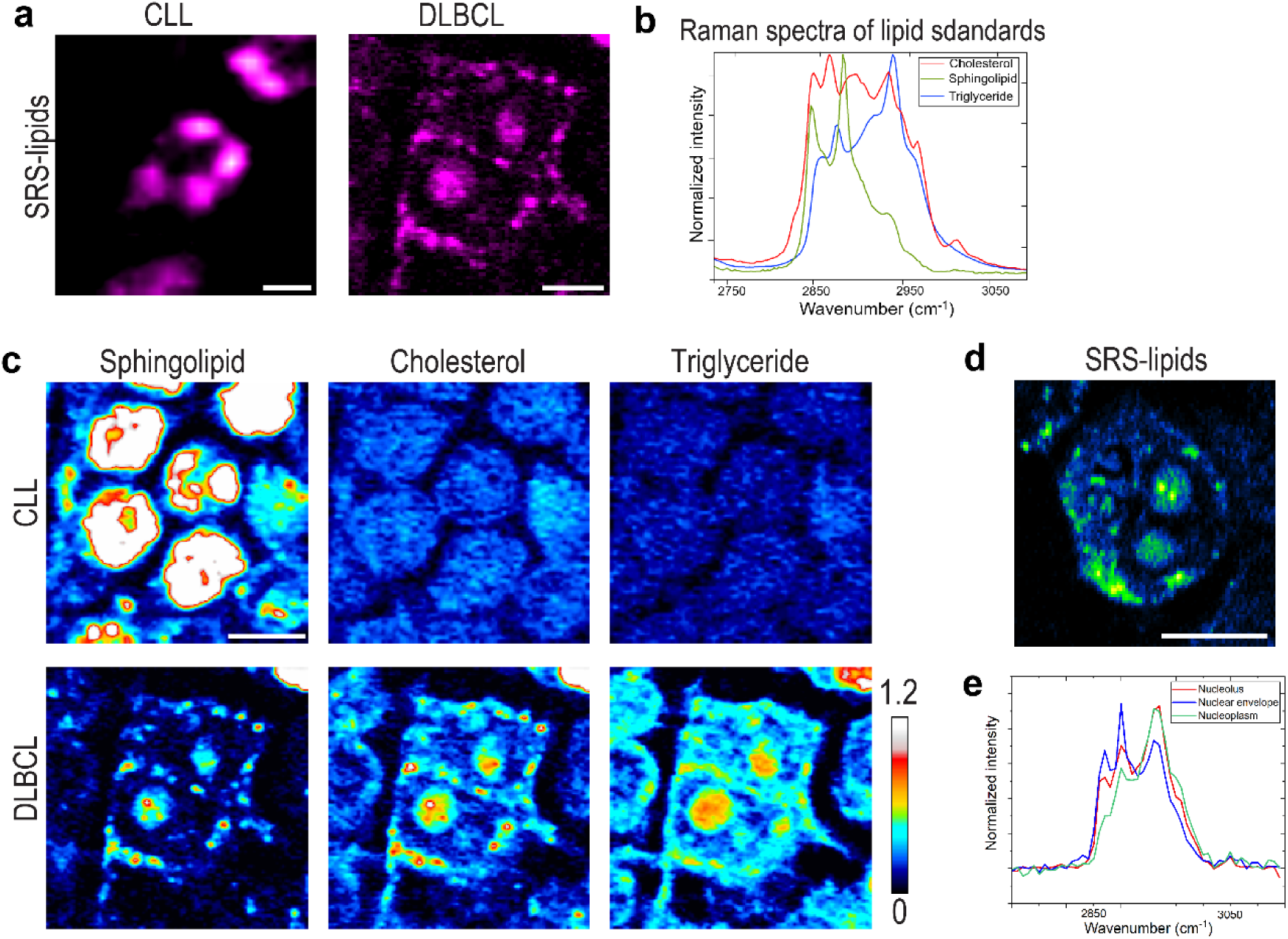
REDCAT reveals lipid metabolic reprogramming in tumor cells. **(a)** High-magnification and super-resolution SRS images highlighting distinct LD morphology in CLL and DLBCL cells. Scale bar, 5 μm. **(b)** Raman spectra of lipid standards, including cholesterol (red), sphingolipids (green), and triglycerides (blue). **(c)** PRM-SRS detection of spatial distribution of cholesterol, sphingolipids, and triglycerides in CLL and DLBCL cells. Scale bar, 10 μm. **(d)** High-magnification and low-exposition SRS image highlighting LDs in nucleolus and perinuclear location of DLBCL cells. Scale bar, 10 μm. **(e)** SRS spectra from different compartments of the nuclei from DLBCL cell.

**Extended Fig. 9.**
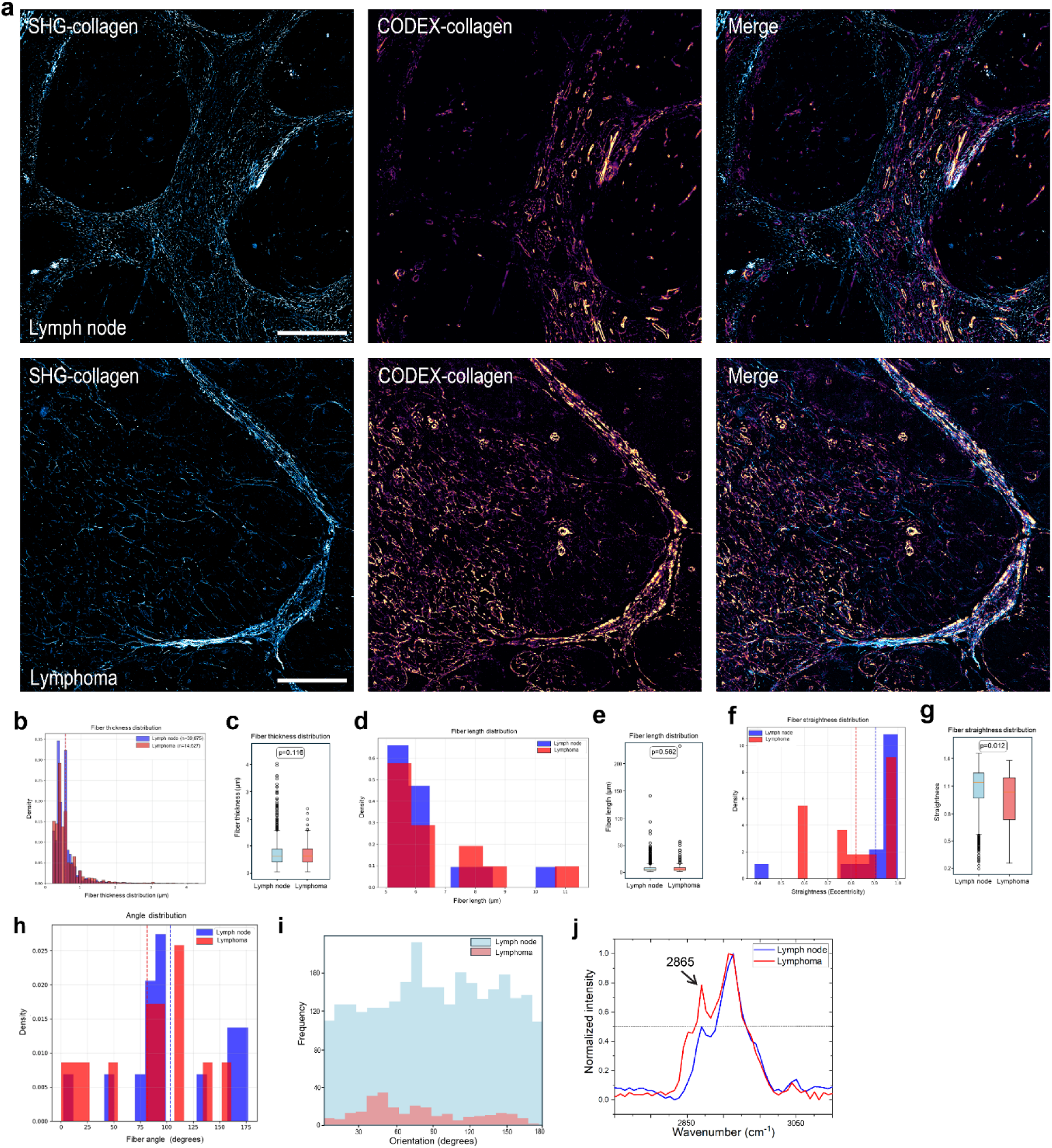
Collagen fiber architecture and composition differences between lymph node and lymphoma. **(a)** Representative SHG images of collagen fibers (type Ⅰ, Ⅱ, Ⅲ) in lymph node (top row) and lymphoma (bottom row) (left); Representative codex images of collagen fibers (type Ⅳ) in lymph node (top row) and lymphoma (bottom row) (middle); merged CODEX and SRS collagen channels (right). Scale bar, 200 μm. **(b, c)** Fiber thickness distribution histograms (b) and boxplots for mean fiber thickness (c) in lymph node and lymphoma. **(d, e)** Fiber length distribution histograms (d) and boxplots (e) comparing lymph node and lymphoma. **(f, g)** Fiber straightness distribution histograms (f) and boxplots (g) between lymph node and lymphoma. **(h)** Angle distribution density plots with mean angle indicated by dashed lines. **(i)** Histogram of collagen fiber orientation shows frequency distribution of angles in lymph node and lymphoma. **(j)** Average SRS spectra of collagen-rich regions from lymph node and lymphoma tissues. The dashed line was to mark the half-height bandwidths of collagen spectra.

**Extended Figure 10.**
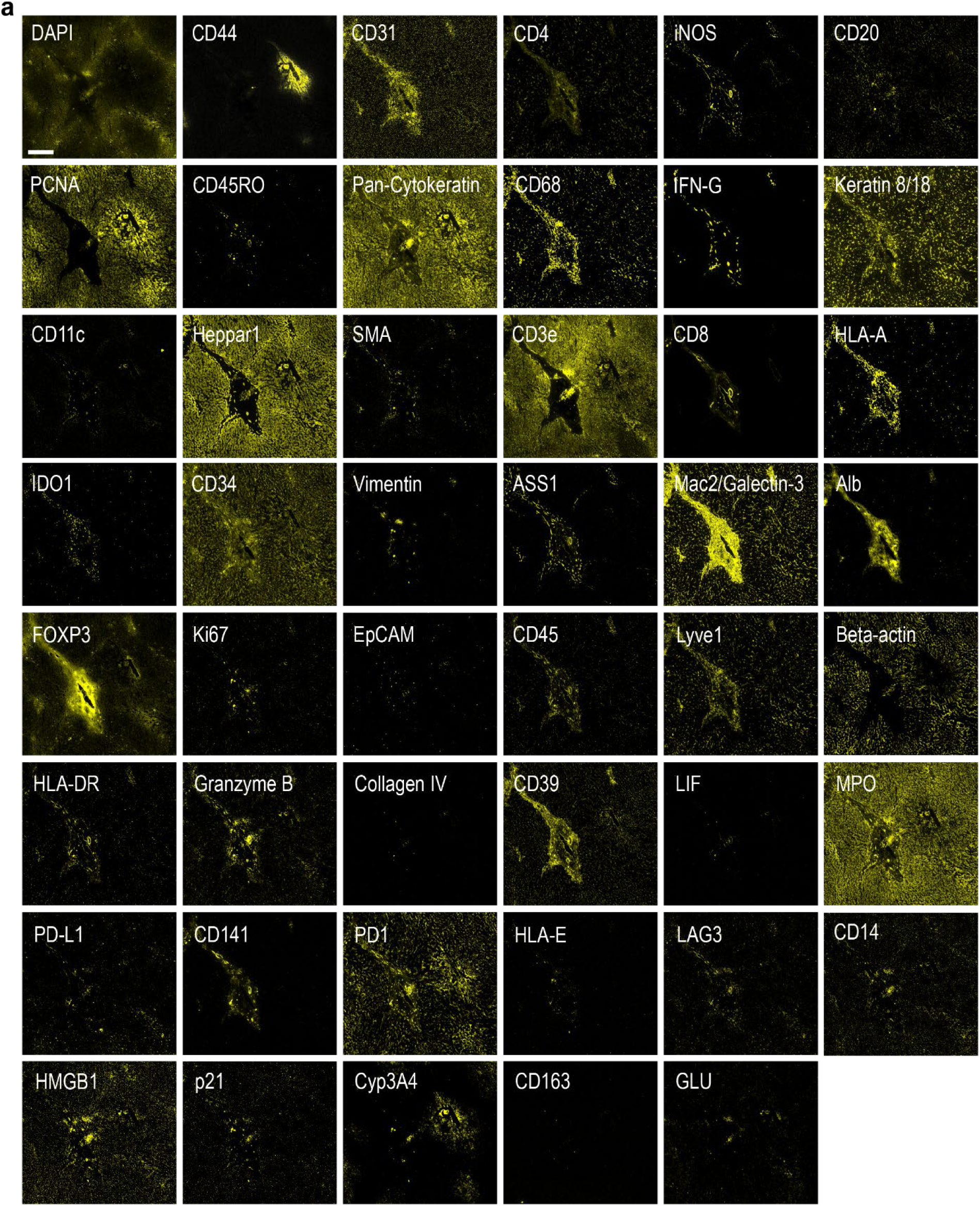
CODEX antibody panel validation for liver tissue. **(a)** Representative single-channel images showing staining patterns for structural, immune, stromal, metabolic, and functional markers in liver tissue. Markers include nuclear and structural proteins (DAPI, PCNA, Beta-actin, Collagen IV, Vimentin), immune lineage markers (e.g., CD3e, CD4, CD8, CD11c, CD14, CD20, CD31, CD44, CD45, CD45RO, CD68, CD141, CD163), hepatic and metabolic enzymes (e.g., HepPar1, ASS1, Cyp3A4, GLUL, Alb), stromal/ECM markers (SMA, Pan-Cytokeratin, Keratin 8/18, Lyve1, Mac2/Galectin-3), functional and activation markers (Ki67, FOXP3, IFN-γ, Granzyme B, LAG3, PD-1, PD-L1, p21, MPO, HMGB1, IDO1, LIF, iNOS), and MHC molecules (HLA-A, HLA-DR, HLA-E). All images were acquired under identical conditions to ensure direct comparison of staining patterns and intensities. Scale bar, 200 μm.

**Extended Figure 11.**
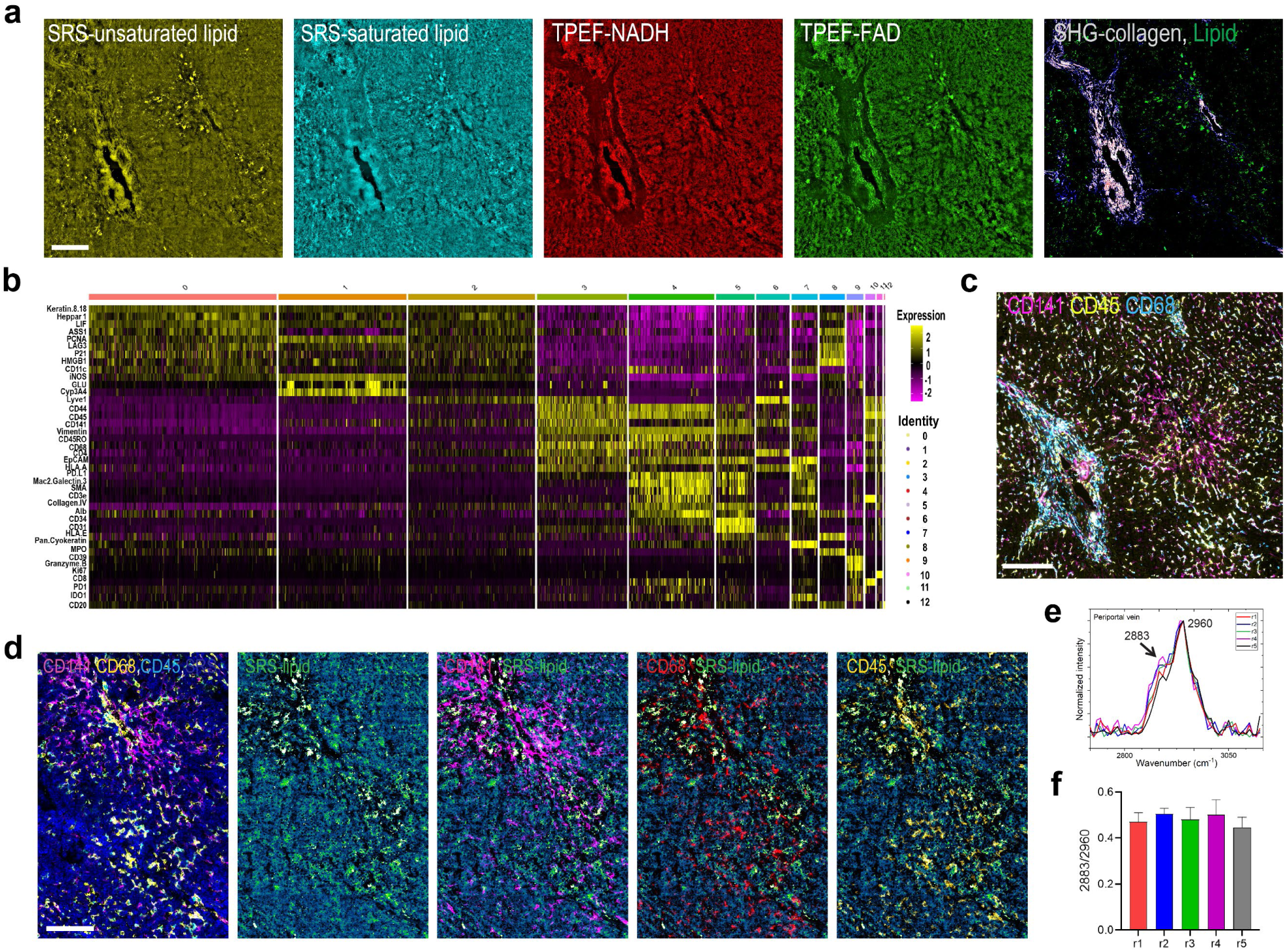
Integrated metabolic and proteomic profiling of liver tissue microenvironments. **(a)** Multimodal imaging channels showing SRS-unsaturated lipid, SRS-saturated lipid, TPEF-NADH autofluorescence, TPEF FAD autofluorescence, and SHG-collagen with overlaid SRS-lipid channel. Scale bar, 200 μm. **(b)** Heatmap of scaled protein marker expression across 12 cell clusters identified by CODEX, with cluster identities indicated above. **(c)** Spatial CODEX map showing localization of CD141 (magenta), CD46 (yellow), and CD68 (cyan) expressing cells in the same ROI. Scale bar, 200 μm. **(d)** CODEX images of another selected region showing overlays between CD141, CD68, and CD45 with SRS-lipid channel (left), and individual SRS lipid channel, and individual CODEX channels for CD141, CD68, and CD45. Scale bar, 200 μm. **(e)** Mean SRS spectra from five representative regions of interest (r1– r5) near PV (indicated in Fig 7l, PV). Concentric regions (r1–r5) extending outward from the PV at 100 µm intervals, marking nuclei within each zone for spatial classification of hepatocytes. SRS spectra were processed by baseline subtraction and smoothing, followed by normalization to peak intensity at 2960 cm^−1^. **(f)** Quantification of the 2883/2960 cm⁻¹ peak ratio across r1–r5 near portal vein, showing no significant variation.

